# Developmental and physiological profiles define drought response diversity and genomic associations in common bean

**DOI:** 10.64898/2026.09.15.751661

**Authors:** Kate Denning-James, Maria Juliana Rodriguez-Cubillos, Jonathan Ashworth, Anthony Hall, Thomas Wood, Caspar C. C. Chater, José J. De Vega

**Affiliations:** Earlham Institute, Norwich Bioscience Institutes, Norwich Research Park, Norwich NR4 7UZ, United Kingdom; Royal Botanic Gardens, Kew, Richmond, Surrey TW9 3AE, United Kingdom; Niab, Crop Improvement, Park Farm Campus, Villa Road, Histon, Cambridge CB24 9AT, United Kingdom Kingdom; Plants, Photosynthesis and Soil, School of Biosciences, University of Sheffield, Sheffield S10 2TN, United Kingdom

**Keywords:** Common bean, *Phaseolus vulgaris*, drought stress, phenotyping, water deficit response, recovery, GWAS

## Abstract

**Background and Aims:** Common bean (*Phaseolus vulgaris* L.) yields are strongly impacted by water deficit, yet there is substantial within-species diversity in how development, biomass accumulation, gas exchange, and photosynthetic performance are coordinated under stress. Based on this variation within a diversity panel, we characterise response profiles, define drought-response strategies, assess how these strategies relate to population structure and gene flow, and identify associated loci.

**Methods:** A panel of 142 common bean accessions representing diverse genetic backgrounds was grown outdoors under controlled water-deficit conditions. Over five weeks, plants were monitored for phenology, biomass, pod production, and leaf traits related to stomatal and photosynthetic performance, including a brief recovery period. Genome-wide association analyses were then performed for developmental, physiological and recovery-related traits.

**Key Results:** Declining soil water availability revealed marked variation among accessions in developmental progression, biomass partitioning, stomatal behaviour and photosynthetic performance. We combined developmental and physiological traits to define above-ground response profiles and classify drought-response strategies in common bean, which were distributed across the diversity panel. GWAS identified multiple QTL and candidate loci associated with developmental, physiological and recovery-related traits.

**Conclusions:** Common bean exhibits extensive diversity in overall above-ground responses to water deficit, likely reflecting local adaptation rather than population structure. Developmental data were essential for differentiating response strategies and, when combined with porometer and fluorometer measurements indicating the level of water stress experienced by the plants, for connecting traits and strategies to genomic variation. These results provide trait relationships, candidate loci and testable hypotheses for validation across environments and for future breeding-oriented studies.

## Introduction

Common bean (*Phaseolus vulgaris* L.) is a globally important legume crop essential for food and nutritional security (Broughton *et al*. 2003). Its high protein and mineral content are especially vital in developing countries, where it often serves as a staple food. Common beans are recognised as a key crop for mitigating climate change due to the ecosystem services they provide, their low environmental footprint, and their role in assuring food and nutritional security (Foyer *et al*. 2016). Agriculture accounts for ∼70% of global freshwater use (Barezzi *et al*. 2024), so climate impacts, such as rising temperatures and irregular rainfall, pose major threats to sustainable yields (IPCC 2023). Common beans are particularly vulnerable to drought, with around 60% of global production exposed to drought during the cropping cycle (Beebe *et al*. 2008; Villordo-Pineda *et al*. 2015). Improving adaptation to water deficit, therefore, remains a priority for bean breeding (Tai *et al*. 2014), but this requires a clearer understanding of how diverse genotypes in the species regulate development and physiology as soil water availability declines.

Plant responses to water deficit are complex, integrating developmental, physiological, and metabolic processes, rather than a single trait in isolation (Tardieu 2012; Araujo *et al*. 2015). In common bean, responses to water deficit include changes in carbon assimilation, transpiration, stomatal conductance, leaf temperature, growth rate, partitioning of assimilates to reproductive organs (inducing leaf senescence and loss), and, under severe stress, flower or seed abortion (Tuberosa 2012; Beebe *et al*. 2013; Polania *et al*. 2020). Because these processes are closely linked, traits such as stomatal conductance, transpiration, chlorophyll-related fluorescence variables, and leaf temperature serve as informative proxies for how plants regulate water loss and maintain photosynthetic function during stress (Smith *et al*. 2019; Marchin *et al*. 2020; Wang *et al*. 2024). Characterising these phenotypes is especially valuable in diversity panels because it enables the identification of contrasting response strategies that frequently coexist within the same crop species.

Drought can refer to different meteorological, agronomic, or physiological phenomena depending on the context (Fioravanti *et al*. 2025). In plant biology, the most relevant factors are the timing, intensity, and duration of reduced water availability relative to plant development (Farooq *et al*. 2017; Habus Jercic *et al*. 2018). In common bean, water limitation during the transition from vegetative growth to flowering or pod set is particularly consequential because reproductive development is highly sensitive to stress (Rosales *et al*. 2012; Labastida *et al*. 2023). Defining the stress scenario precisely is, therefore, essential when comparing genotypes, interpreting physiological data, and linking phenotypes to candidate loci.

Several drought-response strategies have been described in common bean. One is drought escape, in which plants complete their life cycle rapidly before severe stress develops. Others involve different forms of drought avoidance or tolerance expressed through water-use behaviour and canopy maintenance (Polania *et al*. 2022). Drought-escaping accessions are particularly suitable for seasonal, predictable long-term droughts in which rainfall may not return (Shavrukov *et al*. 2017). Drought avoidance strategies include “water spenders” (anisohydric) that accelerate development and “water savers” (isohydric) that delay growth to conserve resources. “Water spenders” sustain photosynthesis during drought stress by keeping their stomata open, even at the cost of water loss. If this strategy allows reproduction and seed set, it can be a valuable crop trait that supports yield stability during prolonged droughts. However, if they cannot complete their life cycle before rains return, this strategy is more appropriate for milder, shorter droughts when growth can resume afterwards (Nesporová *et al*. 2024). On the other hand, “water savers” are plants that reduce photosynthesis by rapidly closing their stomata upon detecting a water deficit and are most effective during milder, shorter droughts; when rainfall returns, stomata reopen, and growth continues (Nesporová *et al*. 2024). A further strategy of interest is functional “stay-green” (SG), in which delayed senescence helps maintain photosynthetic activity under stress (Sofi *et al*. 2021; Borrell *et al*. 2022; Kumar *et al*. 2022; Nunes *et al*. 2022; Polania *et al*. 2022). These strategies are not mutually exclusive, and their value depends on the stress pattern encountered, including its timing, severity and the likelihood of post-stress recovery.

Common bean agrobiodiversity and its natural altitudinal and latitudinal geographic range make it well-suited for studying these differing responses to water availability. The processes of domestication and modern breeding for high-yield varieties have both reduced the genetic diversity of our crops and unintentionally produced varieties with high stomatal conductance and low water-use efficiency (Lei *et al*. 2023; Huang and Zeng 2024). In contrast, locally adapted landraces and admixed backgrounds can retain variation relevant to resilience under water deficit (Renard *et al*. 2023). Common bean has two major domestication gene pools, Andean and Mesoamerican, which have subsequently undergone introgression and regional diversification. This evolutionary history provides a useful framework for assessing whether drought-response traits are associated with broad population structure or recur across distinct genetic backgrounds.

Previous work has identified drought-adapted material in both major domesticated gene pools and related wild relatives, highlighting the importance of traits such as partitioning efficiency, phenology, photosynthetic maintenance, and canopy responses under stress (Beebe *et al*. 2008; Cortés *et al*. 2013; Villordo-Pineda *et al*. 2015; Polania *et al*. 2017; Dramadri *et al*. 2019; Labastida *et al*. 2023). Past studies have also examined different races within these gene pools (Teran and Singh 2002; Munoz-Perea *et al*. 2006; Beebe *et al*. 2008; Polania *et al*. 2020), or included secondary centres of diversification (Cortés *et al*. 2013; Papathanasiou *et al*. 2022), African varieties (Darkwa *et al*. 2016), or admixed accessions (Leitao *et al*. 2021). Much of this research has focused on identifying tolerant lines for cultivar development or testing specific physiological traits in target environments. However, a complementary need is to characterise, across a broad and structured diversity panel, how developmental progression, biomass allocation, gas exchange, and photosynthetic performance combine to produce distinct above-ground response profiles under a specified water-deficit scenario. This involves describing the range of above-ground responses in common bean and investigating whether these strategies are linked to genomic differences, rather than simply declaring specific accessions as “drought tolerant.” This approach is particularly useful because it considers drought response as an integrated, dynamic phenotype, enabling analysis of response diversity, recovery potential, and genomic associations together rather than separately.

Here, we used a diverse panel of common bean accessions from Colombia and neighbouring regions, encompassing Andean, Mesoamerican, and admixed backgrounds, to characterise variation in developmental and physiological responses to a defined water-deficit treatment applied before flowering. Our objectives were: (i) to categorise above-ground quantitative response profiles using phenology, biomass, pod production, transpiration, and photosynthetic traits; (ii) to determine whether integrating developmental progression scoring improves interpretation of physiological measurements under decreasing water availability; (iii) to explore how these response profiles relate to population structure and gene flow between the two domesticated gene pools; and (iv) to identify genomic regions associated with developmental, physiological, and recovery-related traits through GWAS. We thus frame this study as a comparative analysis of response diversity in common bean under controlled stress, validating an evaluation strategy for plant responses that integrates developmental and physiological measurements, and can categorise accessions by their strategies during drought stress. Our results include novel candidate loci for validation in multi-environment and field studies, rather than proposing new types of drought response in common bean, as these have been extensively studied in the species.

## Methods

### Plant materials and genotyping

A diversity panel of 144 common bean (*Phaseolus vulgaris* L.) accessions (Supplementary Table S1) was previously assembled and genotyped, as detailed in (Denning-James *et al*. 2025). These accessions represent a broad range of diversity from Colombia and neighbouring countries, as well as both Andean and Mesoamerican gene pools, and a variety of races. During the study, two accessions (JSPinto and JSBlackwater) died during the water-deficit treatment, and the trial and subsequent downstream analyses were completed with only 142 accessions. The genotypic data from whole-genome resequencing comprised 20.2 million variant loci (∼17.1M SNPs and ∼3.4M indels). This dataset was further filtered for biallelic loci with a minor allele frequency of 1% and thinned to remove clustered SNVs within a 5-bp window using BCFtools v1.12 (Danecek *et al*. 2021). The resulting VCF used for GWAS contained 2,572,124 loci, as detailed in (Denning-James *et al*. 2025). For each accession, seed type (wild, landrace, and heirloom), country of origin, population structure (admixture coefficients estimated by ADMIXTURE; K = 6), seed colour, photoperiod sensitivity, and growth habit information were included from the previous analysis (Denning-James *et al*. 2025).

### Design of the experimental phenotypic evaluation

A water-deficit experiment was performed outdoors at the Niab research station (Histon, UK; 52.246, 0.098) from June to September 2023 (Supplementary Figure S1). The accessions were separated by growth habit (climbing types required support poles) into blocks and organised in a randomised block design with three replicates per accession. All accessions were subjected to water-deficit upon cessation of drip irrigation. Watered controls were included in the design to provide a contrast with the other plants under water-deficit stress. Due to limitations with support poles, only determinate accessions received both treatments (control and water deficit), and growth habit was included in the analyses as appropriate. A 1-metre buffer zone was planted with a commercial common bean variety around the controls to control against water splashing.

As indicated in table 1, all accessions were sown on 19/06/2023 (day 0) in 5-litre pots containing Sylvamix compost (Nitrogen 250 mg/l, Phosphorus 80 mg/l, Potassium 300 mg/l; Melcourt Ltd, UK). Plants were thinned to 1 plant per pot on 06/07/2023 (day 17) after reaching BBCH-scale stage 12, representing the growth of true leaves. The BBCH-scale for beans describes the phenological development of bean plants (Cavalcante *et al*. 2020). During early development, all the plants were irrigated daily, and pots were spaced at ∼20 cm intervals. On 27/07/2023 (day 38), irrigation was stopped to induce the water-deficit treatment. These pots received only water from precipitation thereafter (Supplementary figure S2). The water-deficit treatment was applied after the plants had established and before any of the accessions had reached the flowering stage; on average, BBCH was below 55 on 28/07/2023 (day 39). Control plants received irrigation throughout the trial.

**Table 1:** Time points in the water deficit experiment (June 2023 to September 2023), indicating the stage of the trial and the number of days from sowing.

| Day from sowing | Experimental stage | Date |
| --- | --- | --- |
| 0 | Sowing | 19/06/2023 |
| 17 | Thinning | 06/07/2023 |
| 22 (Week 0) | Normal irrigation | 21/07/2023 |
| 38 | Irrigation removed | 27/07/2023 |
| 39 | Day 1 water deficit | 28/07/2023 |
| 43 (Week 1) | Day 4 water deficit<br>First data recording | 01/08/2023 |
| 46 | Day 7 water deficit | 04/08/2023 |
| 52 (Week 2) | Day 13 water deficit<br>Second data recording | 10/08/2023 |
| 60 (Week 3) | Day 21 water deficit<br>Third data recording | 18/08/2023 |
| 65 (Week 4) | Day 26 water deficit<br>Fourth data recording | 23/08/2023 |
| 67 | Day 1 Recovery | 25/08/2023 |
| 72 (Week 5) | Day 6 Recovery<br>Fifth data recording | 30/08/2023 |
| 79 | Harvesting | 06/09/2023 |
| 82 | Harvesting | 09/09/2023 |

### Environment monitoring

After sowing, fifteen Teros-21 and twelve Teros-12 (METER Group, USA) soil sensors (Cominelli *et al*. 2024) were randomly assigned to pots, in proportion to the number of accessions in each block. These sensors collected hourly data on soil water potential, soil moisture content, soil temperature, and electrical conductivity. ZL6 data loggers (METER Group, USA) were used to record the data. The sensor data were downloaded and analysed in R, and visualised using the ggplot2 package (Wickham 2016).

To examine differences in soil metrics across blocks for each time point, a two-way ANOVA was conducted. Post hoc pairwise comparisons of estimated marginal means were conducted for significant interactions using the emmeans package in R (Lenth 2025). The P-values were adjusted using the Holm method. Data were plotted in R employing ggplot2, paletteer, and ggthemes (Wickham 2016; Hvitfeldt 2021; Arnold 2024).

### Phenotyping data collection and analysis

Leaf water loss and photosynthetic parameters were measured across the diversity panel during the experiment using a handheld LI-600 porometer/fluorometer (LI-COR Environmental, USA). The LI-600 porometer outputs (Table 2) included stomatal conductance to water vapour (*g_sw_*), transpiration (*E*_apparent), leaf vapour pressure deficit (VPDleaf), relative humidity of the sample (rh_s), and leaf temperature (Tleaf), and the fluorometer outputs included electron transport rate (ETR), steady-state fluorescence (Fs), maximum fluorescence (Fm’), and quantum efficiency in light (PhiPS2). Normalised leaf temperature was the difference between Tleaf and the air temperature at data capture. LI-600 measurements were initiated when the water-deficit treatment began (day 38) and were collected weekly between 13:00 and 17:00 BST in random order. For every plant, spot measurements were taken on the central leaflet of fully expanded trifoliate leaves that showed no signs of yellowing or damage.

**Table 2:**
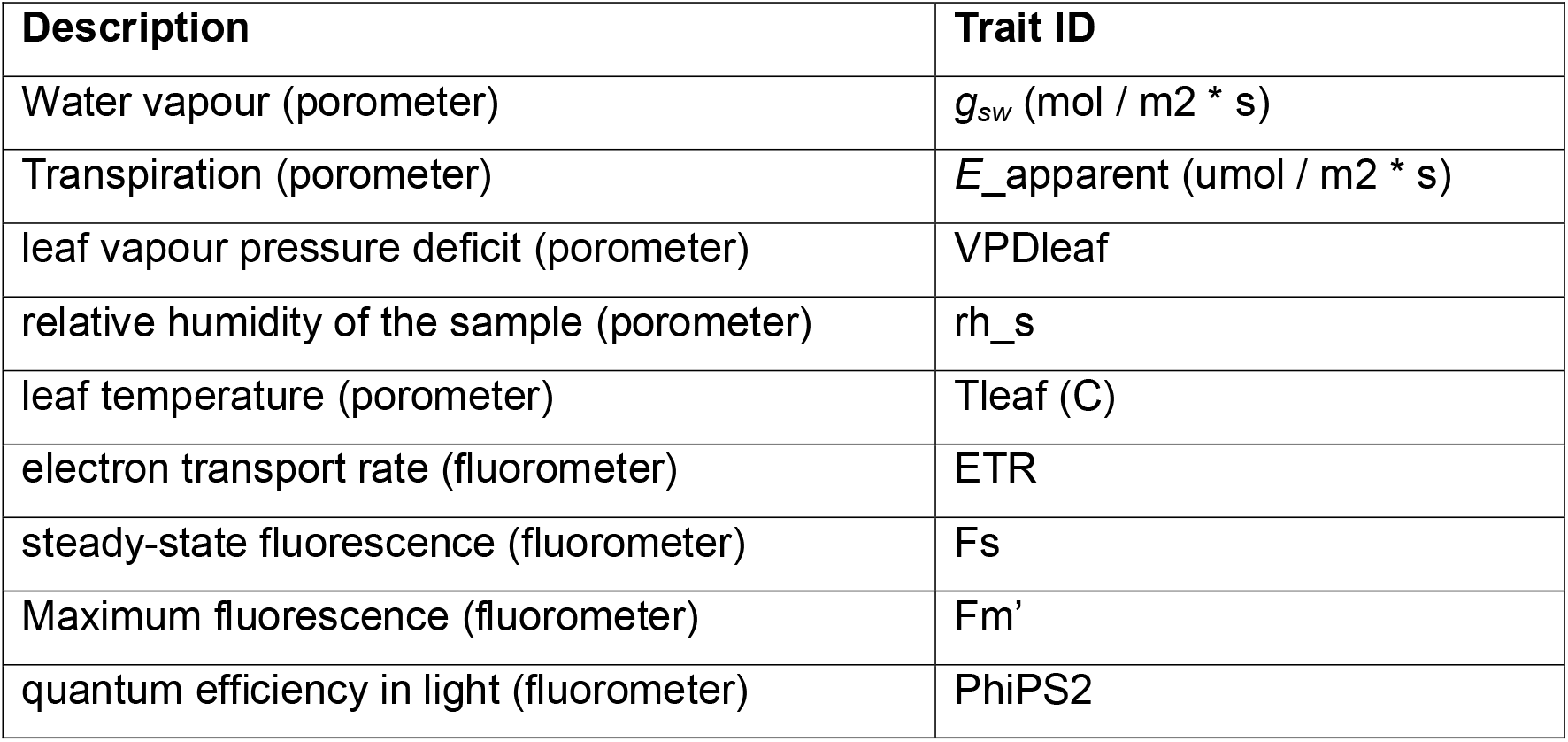
Leaf water loss and photosynthetic parameters measured across the diversity panel during the water-deficit experiment using a handheld LI-600 porometer/fluorometer.

To assess phenological development, BBCH stages were recorded weekly from sowing to harvest (Cavalcante *et al*. 2020). At the end of the experiment (day 79-82), plants were harvested at ground level, their pods counted, and the fresh weight of the pods and leaves measured separately to distinguish reproductive from vegetative growth.

The phenotypic data were tested for normality (p-value > 0.05) using the Shapiro-Wilk test on the residuals with the rstatix package (Kassambara 2023). The statistical analysis of correlations between continuous phenotypic variables was performed in R using Spearman’s rank correlation for non-normal distributions. Differences in distribution between continuous and discrete variables were assessed using the Kruskal-Wallis test for non-parametric data (Kruskal and Wallis 1987; Wei and Simko 2021; Kassambara 2023; R Core Team 2024). The association between two discrete variables was calculated with Cramér’s V and significance with chi-squared (Mangiafico 2025). Results were visualised using the R packages corrplot and ggplot2 (Wickham 2016; Wei and Simko 2021; R Core Team 2024). Analysis of significant differences in growth habits across the porometer and fluorometer variables was completed using a two-way ANOVA, as carried out for the soil monitoring (see previous section).

### Genome wide association studies

A genome-wide association study (GWAS) was conducted using GAPIT v.3 (Wang and Zhang 2021), incorporating three principal components to account for population structure across all phenotypes collected (Supplementary Table S1). Phenotype scores obtained in weeks 1 and 2 were labelled as “initial response to water deficit”, while the difference between weeks 4 and 5 was labelled as “recovery response”. To estimate the response’s magnitude, we also calculated the differences between scores from weeks 1 and 4 (“1 to 4”), and weeks 1 and 5 (“1 to 5”), along with their averages across all dates, the total sum of all dates, and the scaled sum (normalised scores). Harvest data included pod weight, foliar weight, and partitioning weight, i.e., the ratio of pod weight to foliar weight (Polania *et al*. 2016; Rehling *et al*. 2021). Significance of the new scores was assessed using the Wilcoxon signed-rank test for non-parametric paired data (Rosner *et al*. 2006).

GAPIT was run on the entire diversity panel using the models “Bayesian-information and Linkage-disequilibrium Iteratively Nested Keyway” (BLINK) (Huang *et al*. 2019), “Fixed and Random Model Circulating Probability Unification” (FarmCPU) (Liu *et al*. 2016), “Multiple Locus Mixed Linear Model” (MLMM) (Segura *et al*. 2012), and “Mixed Linear Model” (MLM) (Zhang *et al*. 2010). BLINK, FarmCPU, and MLMM were selected as multi-locus models for different heritability levels to enhance statistical power (Huang *et al*. 2019; Merrick *et al*. 2022; Cebeci *et al*. 2023). MLM was used as a single-locus analysis for baseline comparison with the other models. When using the BLINK model, GAPIT was run with the parameter “Random.model=TRUE,” so Phenotypic Variance Explained (PEV) values were not calculated. To assess how well the models fit the phenotypic data, quantile-quantile (QQ) plots were employed. All plots were generated in R using the ‘ggplot2’ package (Wickham 2016).

### QTL identification and candidate gene prediction

Marker trait associations (MTAs) were investigated based on significance (-log10(p-value) > 7) and when the MTA was identified with two different phenotypic traits across any model. QTLs were defined as ±100 kbp from the MTA; this is an approximation based on an LD decay of approx. 114 kb at R2 = 0.25 for this panel, as reported and discussed in (Denning-James *et al*. 2025).

To prioritise causative genes within QTLs, the criteria previously tested in (Denning-James *et al*. 2025) were followed: the Andean reference genome (*Phaseolus vulgaris* G19833 v2.1 (Schmutz *et al*. 2014; Diesh *et al*. 2023) was visualised in JBrowse and used to identify genes within the QTLs. SnpEff was then used to identify non-synonymous “high impact” mutations (Cingolani *et al*. 2012). eggNOG-mapper v2 provided functional annotations for genes in the QTLs based on orthologue assignments (Cantalapiedra *et al*. 2021). Genes annotated with GO terms GO:0009819, GO:2000070, GO:1902584, GO:0009414, GO:0009415, GO:0042630, GO:0042631, GO:0097207, GO:0006970, GO:0009651, GO:0009269, GO:0009992, or GO:0080148 were labelled as related to drought stress and/or abiotic stress responses, and prioritised.

Selected candidate genes were further investigated using PhytoMine (Goodstein *et al*. 2012), which includes the reference *Phaseolus vulgaris* v.2, homology analyses against the non-redundant (nr) protein database at NCBI, and, if no gene function could be identified in closer relatives, comparisons against the TAIR database (Huala *et al*. 2001). The loci were compared to previous studies and literature using PulseDB. QTLs and markers were mapped to the reference genome to estimate the conversion from cM to Mb and were visualised in JBrowse (Humann et al. 2019).

## Results

### Inducing water-deficit stress in the diversity panel

An experiment to induce water-deficit stress was conducted outdoors, in which all accessions were sown (day 0) and, once drip irrigation was stopped on day 38, subsequently subjected to water-deficit stress. Throughout the experiment, the average day length was 15.4 hours, and the ambient temperature from sowing to harvesting was mild and consistent, averaging 17.6 °C, with maximum and minimum temperatures of 24 °C and 13 °C, respectively (Supplementary figure S2). The mean precipitation between the time when irrigation stopped (day 38) and harvest (day 79) was 1.73 mm/day (Supplementary figure S2).

Soil water content (m^3^/m^3^) quickly reflected the treatment (figure 1): it decreased from over 0.2 m^3^/m^3^ (day 38) to below 0.1 m^3^/m^3^ within less than 4 days after irrigation was stopped. Differences between the control and stressed accessions were already significant 7 days after irrigation was stopped (week 1). Soil water content gradually decreased, reaching undetectable values (recorded as 0 m^3^/m^3^) by week 3 in nearly all treated accessions and by week 4 in all accessions except controls (figure 1). However, precipitation (rainfall) increased after the measurements in week 4 (supplementary figure S2), leading to an increase in soil water content, as plants were not sheltered from rainfall. In week 5, soil water content (m^3^/m^3^) values were equivalent to those observed in week 3. This precipitation was insufficient to fully offset the water previously lost, as supported by the continued decline in soil metrics at week 5 (Supplementary figure S3), but it was sufficient to induce a “recovery response” in the treated plants (presented in the following results section).

**Figure 1.**
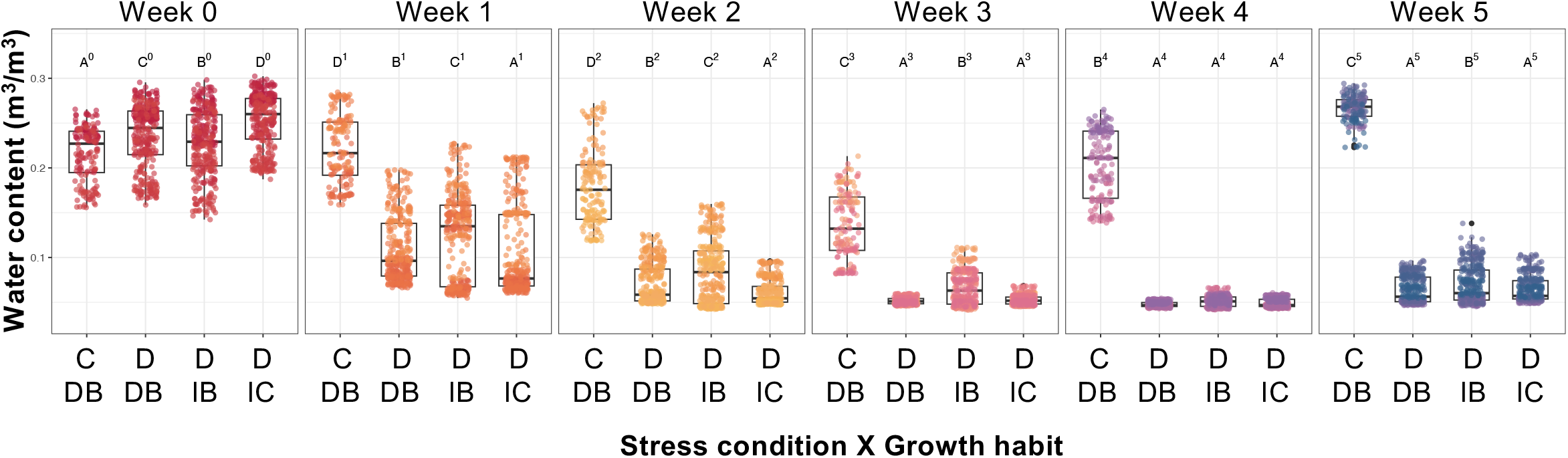
Soil water content declines following the imposition of water deficit. Soil water content (m³/m³) measured from week 0 to week 5. Points are individual sensor measurements, and boxplots summarise distributions. Groups are C DB, irrigated control determinate bush; D DB, water-deficit determinate bush; D IB, water-deficit indeterminate bush; and D IC, water-deficit indeterminate climbing. Colours indicate sampling week. Different letters indicate significant differences within each time point at α = 0.05.

Soil matric potential (kPa) and bulk electrical conductivity (Bulk EC; mS/cm) showed patterns similar to those of soil water content, with measured values significantly different between control and treated plants from week 1 onward, peaking at weeks 3 and 4 (supplementary figure 3). Soil temperature ranged from 17.2°C to 20.8°C across weeks and did not differ significantly between the control and water-deficit groups (supplementary figure 3).

### Measuring quantitative responses to water-deficit stress in the diversity panel

Porometer and fluorometer data (*E*, *g_sw_*, rh_s, VPDleaf, Tleaf, PhiPS2, ETR, Fm, Fs) were collected weekly for all accessions under water-deficit stress and irrigated control conditions. There were significant correlations (p<0.01) between most of the captured stomatal and photosynthetic parameters (figure 2). Stomatal conductance (*g_sw_*), Relative humidity (RH), electron transport rate (ETR), steady-state chlorophyll fluorescence (Fs), transpiration (*E*_apparent), maximum fluorescence yield (Fm’) and effective quantum yield of PSII (PhiPS2) were all strongly positively correlated to each other (r>0.6). Leaf temperature (Tleaf) and leaf vapour pressure deficit (VPDleaf) were strongly positively correlated with each other (r=0.98), but they were negatively correlated with the previous group of measurements (*g_sw_*, RH, etc.).

**Figure 2.**
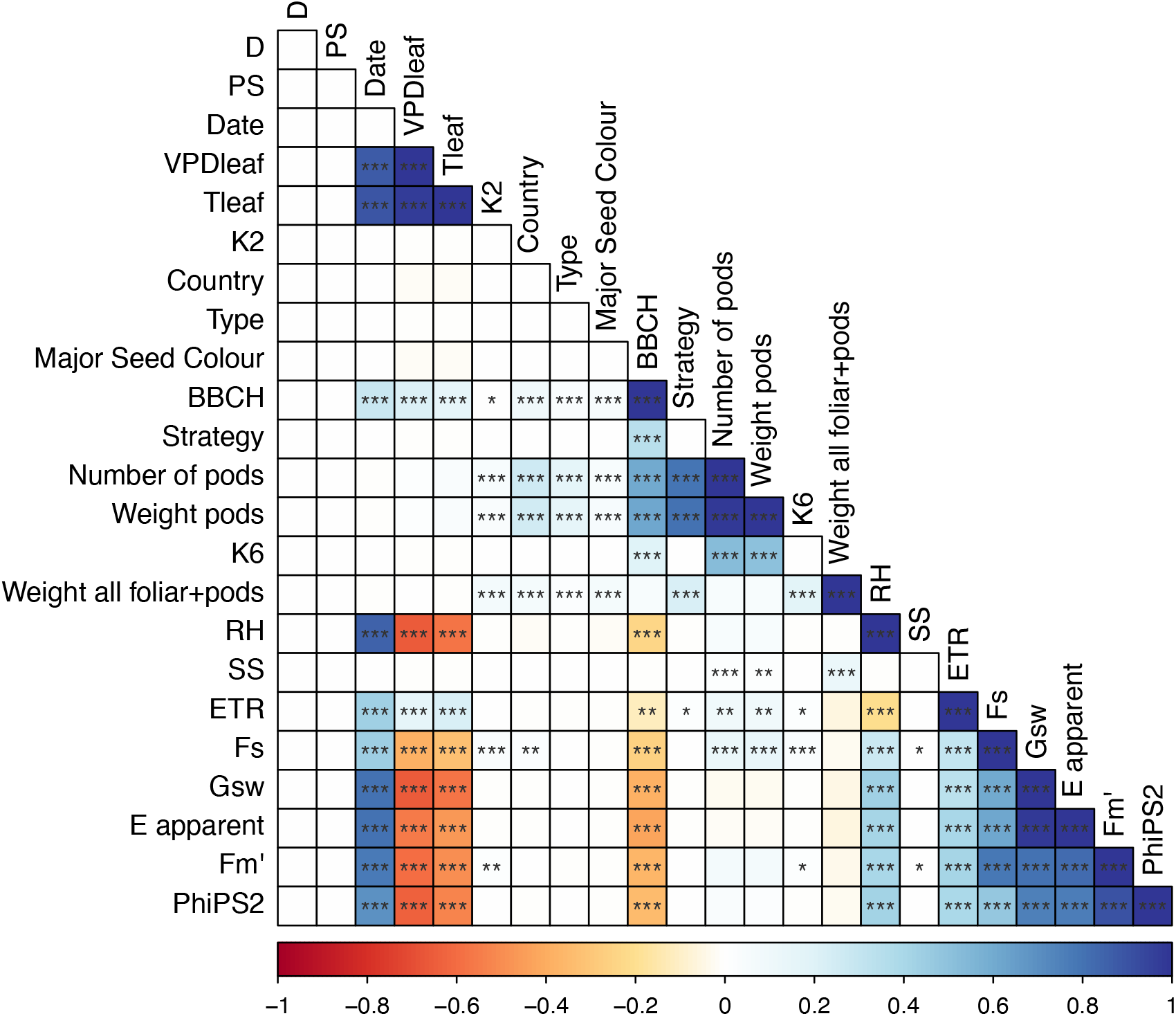
Developmental, yield and physiological traits show coordinated responses to water deficit. Association matrix for developmental, physiological, yield and accession-level variables in 142 water-deficit-treated accessions. Cell colour indicates association direction and magnitude, from negative (red) to positive (blue). Asterisks indicate significance (\**P* < 0.05, \*\**P* < 0.01, \*\*\**P* < 0.001). D, determinacy; PS, photoperiod sensitivity; VPDleaf, leaf vapour-pressure deficit; Tleaf, leaf temperature; K2 and K6, ADMIXTURE population structure; RH, relative humidity; SS, seed size; ETR, electron transport rate; Fs, steady-state fluorescence; gsw, stomatal conductance; E, apparent transpiration; Fm′, maximum fluorescence yield; PhiPS2, ΦPSII.

Accounting for redundancy among highly correlated variables, we focused on the changes in stomatal conductance (Gsw, *g_sw_*) and leaf T (Tleaf) across weeks (figure 3). As expected, there were no differences between control and treated plants before treatment (week 0). After 3 weeks of treatment (figure 3A), *g_sw_* was nearly 0 mol m⁻² s⁻¹ in stressed accessions, and Tleaf was higher (over 25°C) in treated accessions than in the irrigated controls (under 25°C). These observations were consistent across replicates, times, and treatments for each accession’s *g_sw_*and Tleaf values (supplementary figures S4 and S5). There were no significant differences in porometer/fluorometer measurements among growth habits, with similar values observed in water-deficit-treated determinate and indeterminate accessions (Figures 3B and 3C; and supplementary Figure S6).

**Figure 3.**
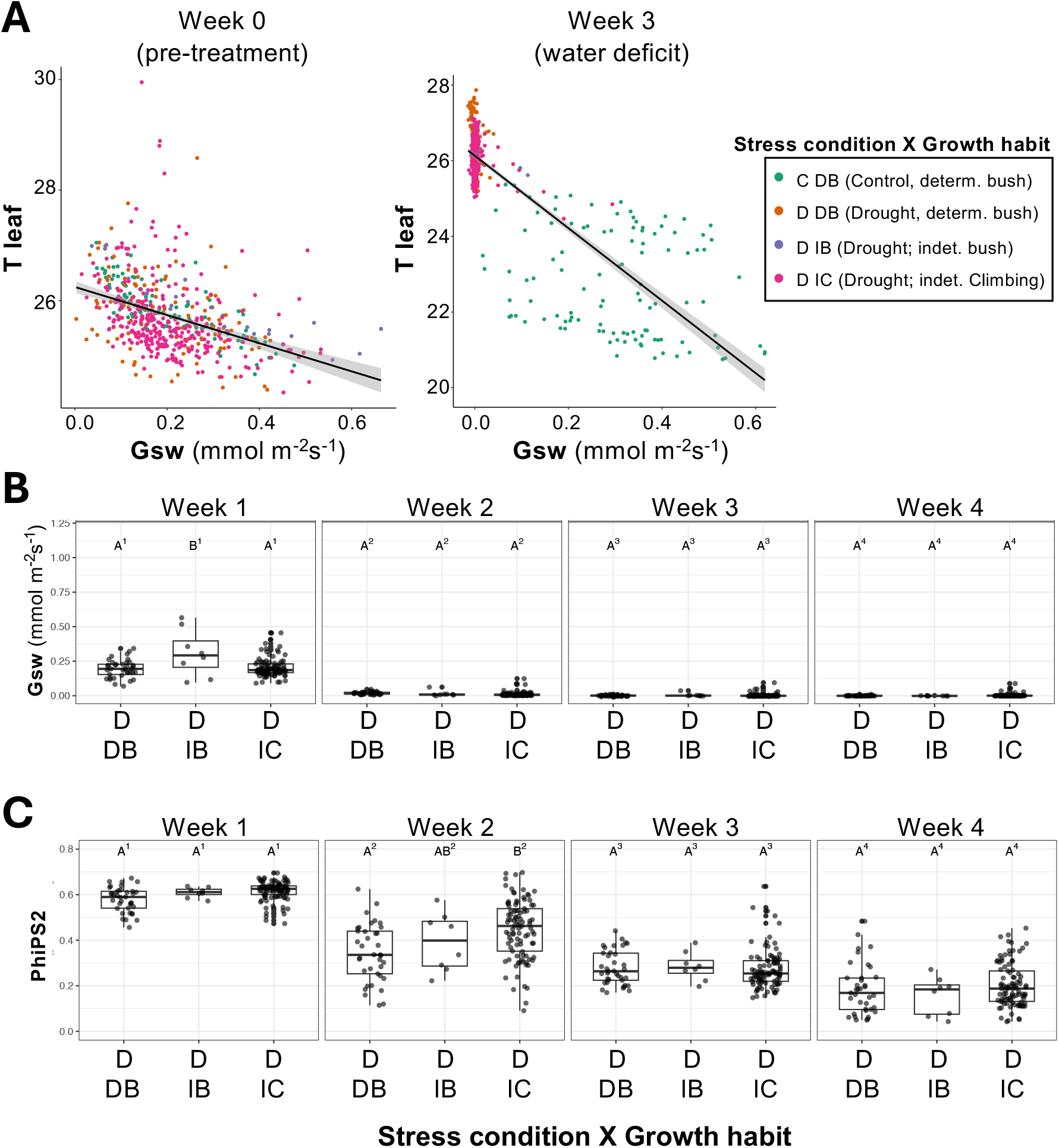
Stomatal and photosynthetic responses track increasing water deficit across growth habits. (A) Relationship between leaf temperature (Tleaf) and stomatal conductance (gsw) before treatment and during water deficit. Points are individual measurements; colours indicate treatment × growth habit; black lines show fitted relationships with confidence intervals. (B, C) Variation across weeks 1–4 in (B) Gsw and (C) ΦPSII. Boxplots summarise distributions. C DB, control determinate bush; D DB, water-deficit determinate bush; D IB, water-deficit indeterminate bush; D IC, water-deficit indeterminate climbing. Different letters indicate significant differences at α = 0.05.

Additionally, phenological development was measured, including harvest information (fresh foliar weight, pod weight, and number) and developmental scoring (BBCH and photoperiod insensitivity). There were significant differences between control and treated plants from week 2; all control plants increased their BBCH and progressed normally through development (Figure 4A). Some accessions under water-deficit treatment reached BBCH values similar to those of the controls, i.e., showing no apparent developmental penalty from the stress. On the contrary, most accessions under water-deficit treatment did not increase their BBCH scores between weeks 2 and 4, i.e., paused phenological development (Figure 4A). Pod number and pod weight were highly correlated (r = 0.99) but not with foliar weight, indicating a trade-off between these traits. Exploring these further (Figure 4B), water deficiency had an obvious effect on both foliar and pod weight, with the top-performing accessions under water deficit reaching the same weights (∼50 g total pod weight) as the worst-performing control ones.

**Figure 4.**
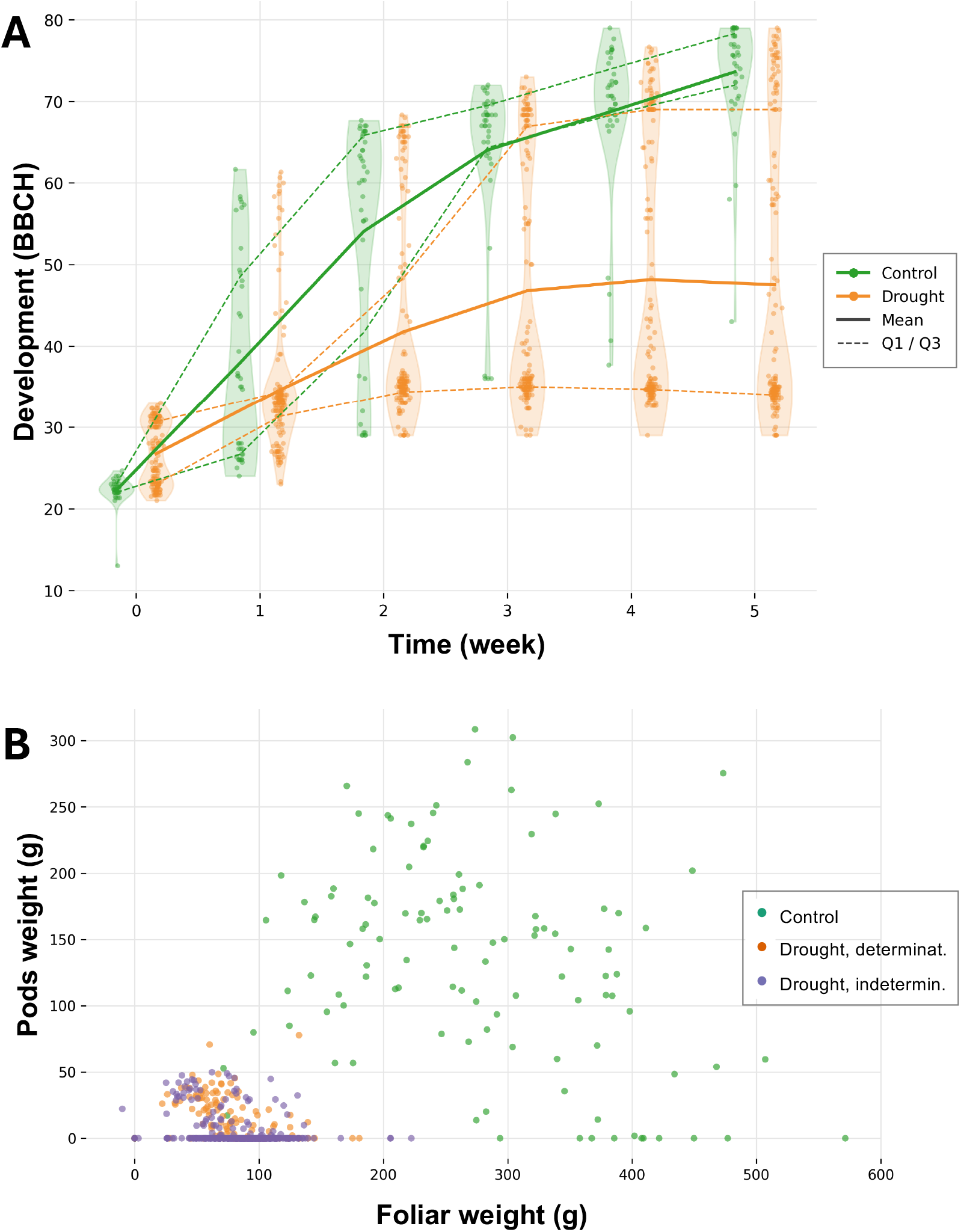
Water deficit alters developmental progression and biomass allocation in common bean. (A) Developmental progression scored using BBCH across five weeks. (B) Relationship between pod fresh weight and foliar fresh weight at harvest, grouped by stress condition and growth habit.

BBCH correlated positively with pod weight (r=0.62) and number of pods (r=0.61), as higher BBCH scores correspond to the development and ripening of fruits (Figure 2). However, BBCH did not correlate with foliar weight. BBCH and harvest traits negatively correlated with *g_sw_*(r=-0.39) and its correlated traits (RH, ETR, Fs, etc.), likely confounded by time and developmental stage.

There were significant differences (p<0.001) in harvested pod weight and number of pods between determinate and indeterminate water-deficit accessions (supplementary figures S7A and 7D). This significant difference between the two groups was not observed in foliar weight or total aerial weight at harvest (supplementary figures S7B and S7C).

### Water-deficit response strategies in common bean

As previously noted (figure 4), the phenological response to water-deficit stress varied significantly among accessions, with some progressing through phenological stages similarly to controls, while others, by contrast, ceased flowering and pod production entirely. By analysing the quantitative phenotypic data collected, we classified these response strategies as “stay-green”, “saver”, “spender”, “prioritised yield”, “drought susceptible”, and “nominal growth” (figure 5). An additional label was used for “no classification” (Table 3). All these categories contained members from both the Andean and Mesoamerican gene pool, except “stay-green”, which only included Andean accessions in our panel.

**Figure 5.**
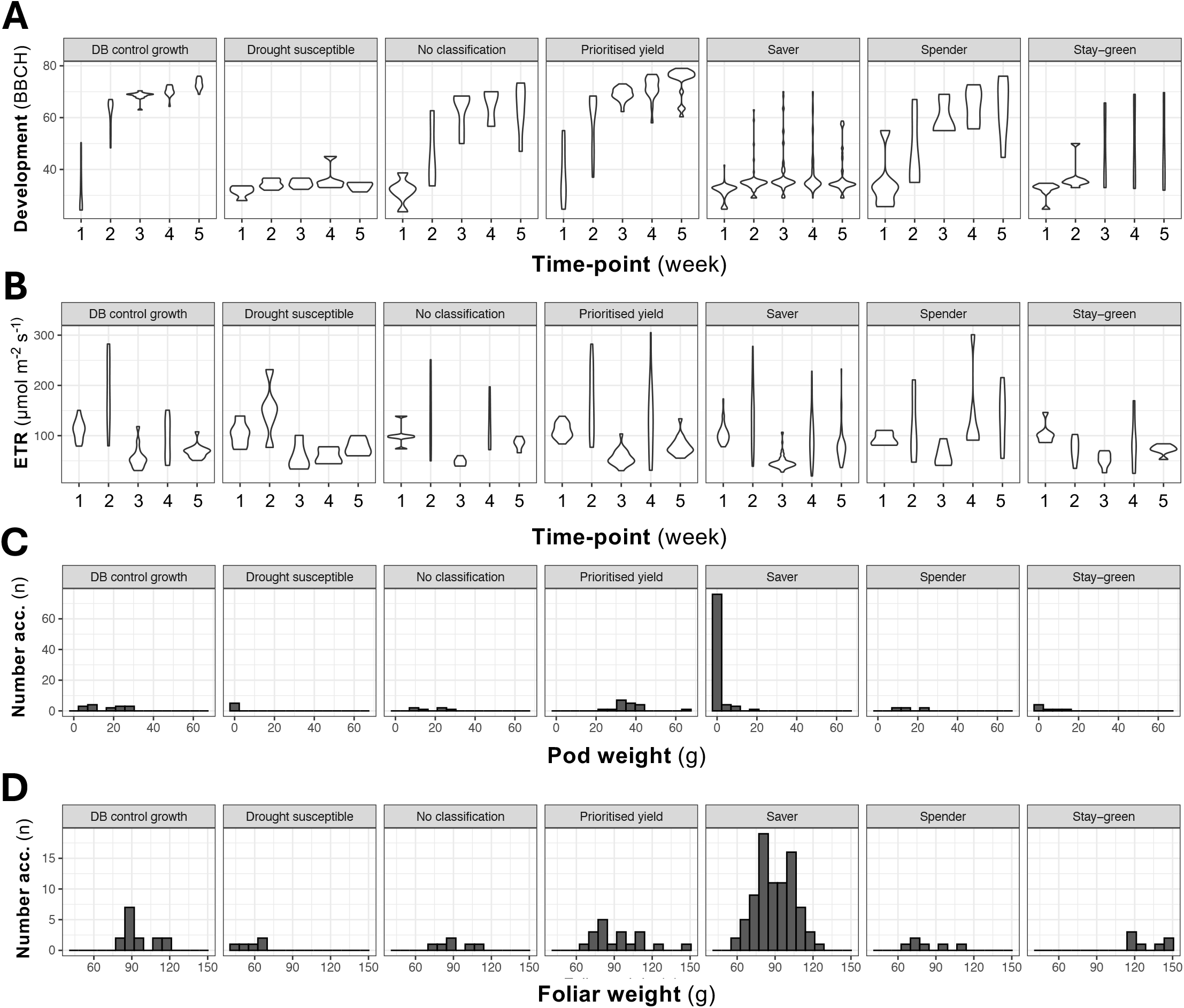
Drought-response strategies differ in development, photosynthetic performance and biomass allocation. (A) BBCH progression and (B) electron transport rate (ETR) across five weeks. (C) Pod fresh weight and (D) foliar fresh weight distributions among drought-response strategy classes.

**Table 3:**
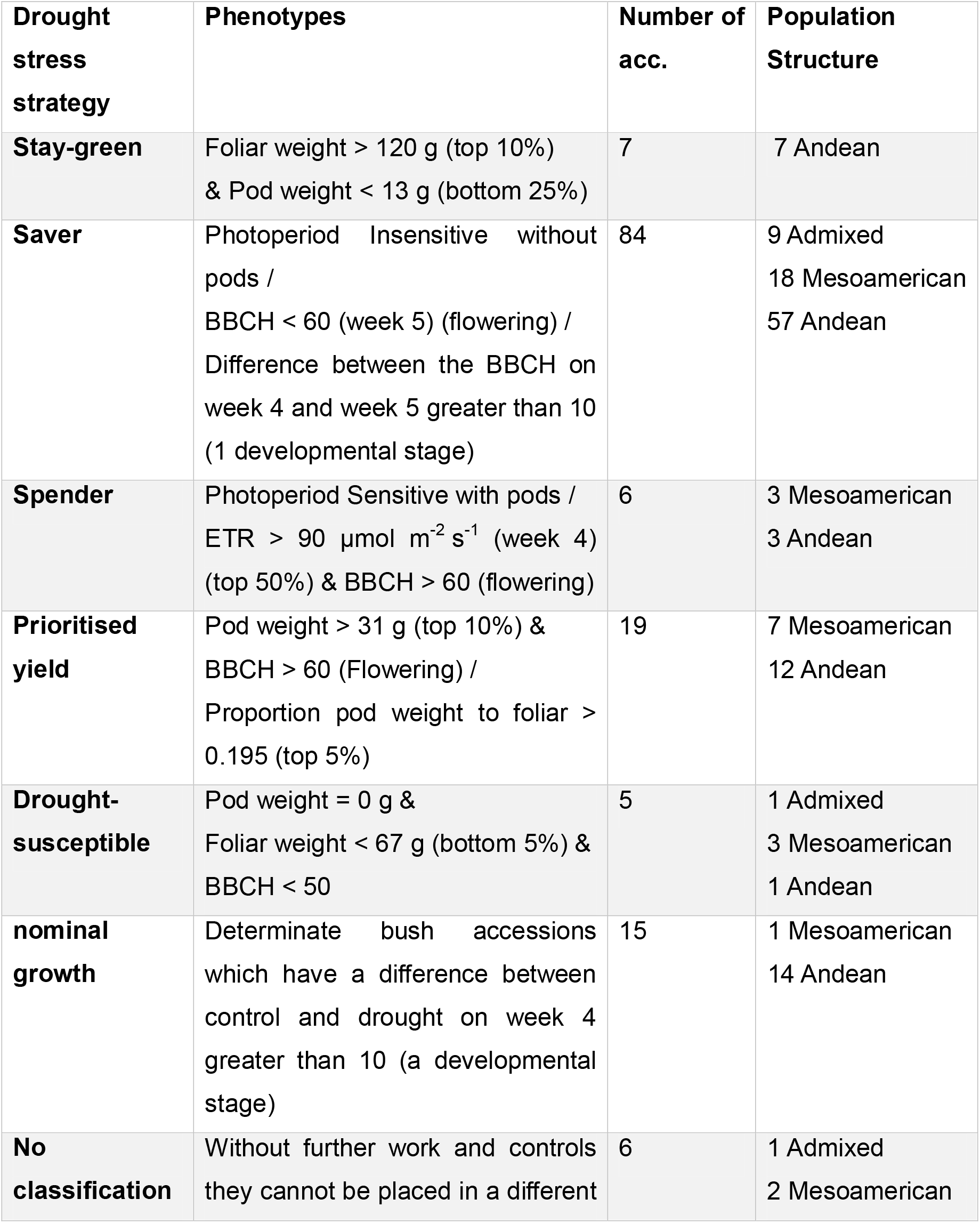
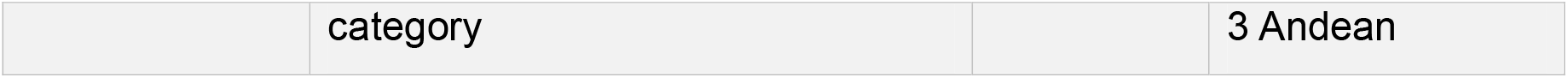
Classification into drought stress strategies based on phenotypic characteristics, photoperiod sensitivity and growth habit data. Data collected at week 4 (four weeks after irrigation was stopped) were selected for this analysis, as this time point reflected the maximum duration of water deficit.

| **Drought stress strategy** | **Phenotypes** | **Number of acc.** | **Population Structure** |
| --- | --- | --- | --- |
| **Stay-green** | Foliar weight > 120 g (top 10%) & Pod weight < 13 g (bottom 25%) | 7 | 7 Andean |
| **Saver** | Photoperiod Insensitive without pods /<br>BBCH < 60 (week 5) (flowering) /<br>Difference between the BBCH on week 4 and week 5 greater than 10 (1 developmental stage) | 84 | 9 Admixed<br>18 Mesoamerican<br>57 Andean |
| **Spender** | Photoperiod Sensitive with pods /<br>ETR > 90 $\mu\text{mol m}^{-2} \text{s}^{-1}$ (week 4) (top 50%) & BBCH > 60 (flowering) | 6 | 3 Mesoamerican<br>3 Andean |
| **Prioritised yield** | Pod weight > 31 g (top 10%) & BBCH > 60 (Flowering) /<br>Proportion pod weight to foliar > 0.195 (top 5%) | 19 | 7 Mesoamerican<br>12 Andean |
| **Drought-susceptible** | Pod weight = 0 g & Foliar weight < 67 g (bottom 5%) & BBCH < 50 | 5 | 1 Admixed<br>3 Mesoamerican<br>1 Andean |
| **nominal growth** | Determinate bush accessions which have a difference between control and drought on week 4 greater than 10 (a developmental stage) | 15 | 1 Mesoamerican<br>14 Andean |
| **No classification** | Without further work and controls they cannot be placed in a different | 6 | 1 Admixed<br>2 Mesoamerican |
|  | category |  | 3 Andean |

“Stay-green” accessions had a pod weight in the lowest quartile (below 13 g in this experiment; figure 5C) and a high foliar weight in the top 10% (over 100 g in our experiment; figure 5D). “Saver” accessions were photoperiod-insensitive accessions, that were expected to produce pods, but did not produce pods (or produced small, aborted pods) under water deficit stress (figure 5C). Regarding phenotype scores, this group did not reach flowering (BBCH below 60; figure 5A) or aborted flowering in the last weeks (negative BBCH between weeks 4 and 5; figure 5A). “Spender” accessions were photoperiod-sensitive plants that produced flowers and pods (BBCH > 60; figure 5A). They also showed ETR (electron transport rate) values greater than 90 (an indicator of relatively high photosynthetic electron transport) in week 4, after the longest period of water deficit (figure 5B). Accessions labelled as “prioritised yield” reached flowering normally (BBCH >60; figure 5A) and had a pod weight in the top 10% (figure 5C), or in some cases better evidenced by a ratio between pod weight and foliar weight in the top 5% (high pod to low foliar weight). “Drought susceptible” accessions did not progress beyond budding and produced no pods (Figure 5C), as evidenced by foliar weight in the bottom 5% and a BBCH below 50 (Figure 5A). “Nominal growth” included treated accessions that had similar growth and development, i.e., were in the same overall developmental stage as their respective controls (BBCH difference < ±10). Finally, any accessions which did not fit into these categories were labelled as “no classification”.

Significant associations (figure 2) were determined between tolerance strategies and estimated seed size (ES = 0.3), Type (Wild, Landrace, commercial, Heirloom; ES = 0.3), Country of origin (ES = 0.3), Major seed colour (ES = 0.34), K6 population structure (‘Admixed K6’, ES = 0.35) and growth habit (GH, ES= 0.5).

### Induced “recovery response” in the treated plants

In week 5, soil water content (m^3^/m^3^) “reverted” to values equivalent to those observed in week 3 because of increased rainfall. This precipitation was insufficient to fully offset the water previously lost, as shown by the continued decline in soil metrics (Figure 1A, supplementary figure 3), but it was sufficient to induce a “recovery response” in the treated plants. To quantify the accessions’ “recovery response”, we quantified the difference between week 4 and 5 across multiple traits (Supplementary figure S8). Significant differences (p<0.001) were identified for the electron transport rate (ETR; 92 to 74 μmol/m^2^ s), leaf temperature (Tleaf; 32.4°C to 20.5°C), transpiration (*E*_apparent; 0.008 to 0.8 mmol/m^2^/s), quantum yield of Photosystem II (PhiPS2; 0.2 to 0.5), Fs (steady-state chlorophyll fluorescence; 97 to 122), relative humidity (rh_s; 30% to 46%), *g_sw_* (stomatal conductance; 0.0002 to 0.07 mol/m^2^ s) and Fm’ (maximum fluorescence yield; 119 to 259).

### GWAS on water-deficit response traits and strategies

A GWAS was performed with all the phenotypes scored during the trial (Supplementary table S2) using the models BLINK, FarmCPU, MLM and MLMM, and MTAs were considered significant when-log10(p-value) > 7 (Figure 6). A total of 36 QTLs were then selected for having significant association with more than one phenotype (Table 4). The QQ plots were analysed to determine the suitability of the model fit (Supplementary figure S9).

**Figure 6.**
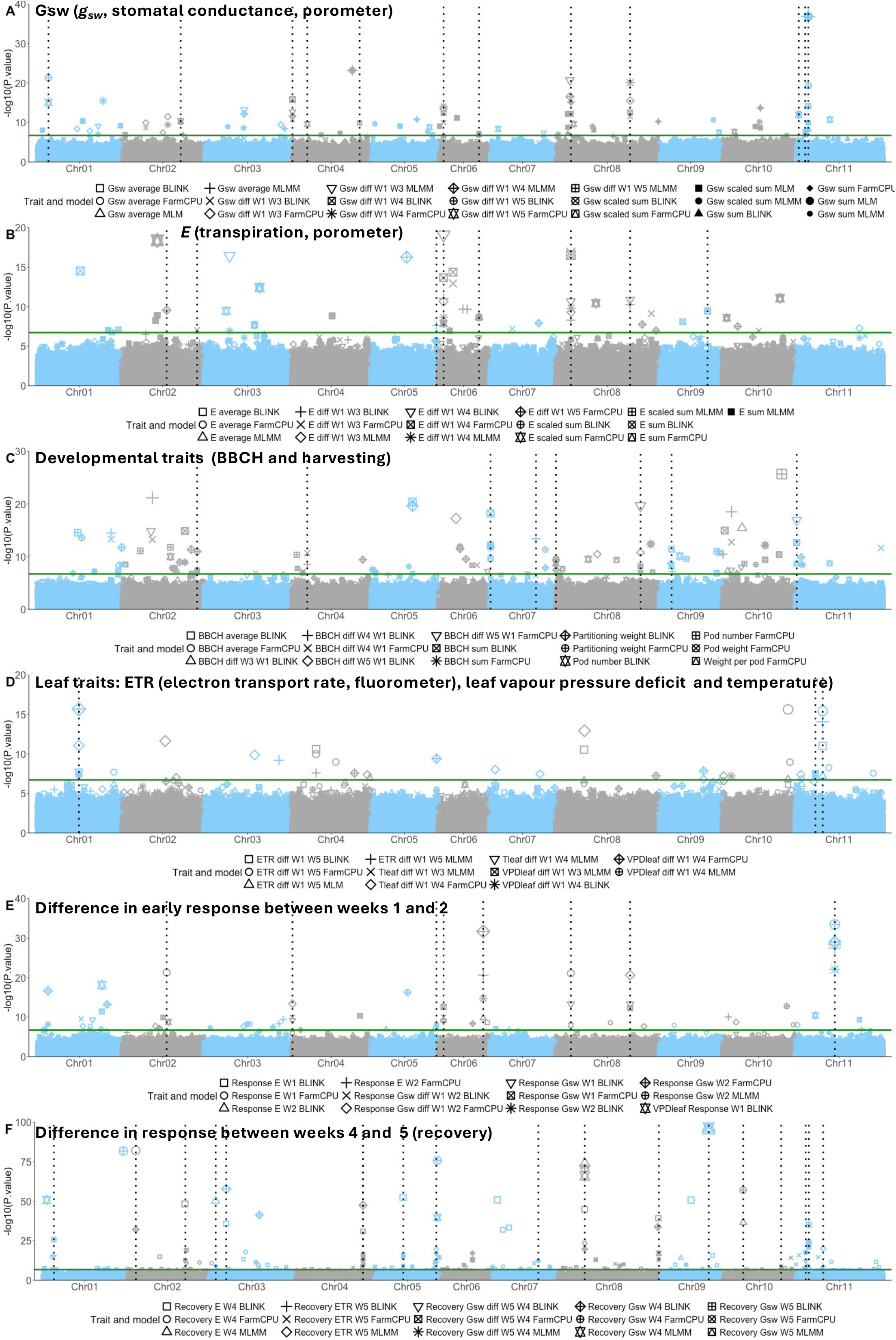
Genome-wide association analyses identify loci associated with developmental, physiological and recovery responses to water deficit. Manhattan plots for (A) stomatal conductance (gsw), (B) transpiration (E), (C) developmental and harvest traits, (D) ETR, VPDleaf and Tleaf, (E) early responses between weeks 1 and 2, and (F) recovery responses between weeks 4 and 5. GWAS were performed on 142 accessions using BLINK, FarmCPU, MLMM and MLM. The x-axis shows genomic position and the y-axis −log₁₀(*P*). Alternating colours distinguish chromosomes, symbol shapes indicate phenotype/model combinations, the horizontal green line marks the significance threshold (−log₁₀*P* = 7), and vertical dotted lines indicate QTLs associated with at least two phenotypes.

**Table 4:**
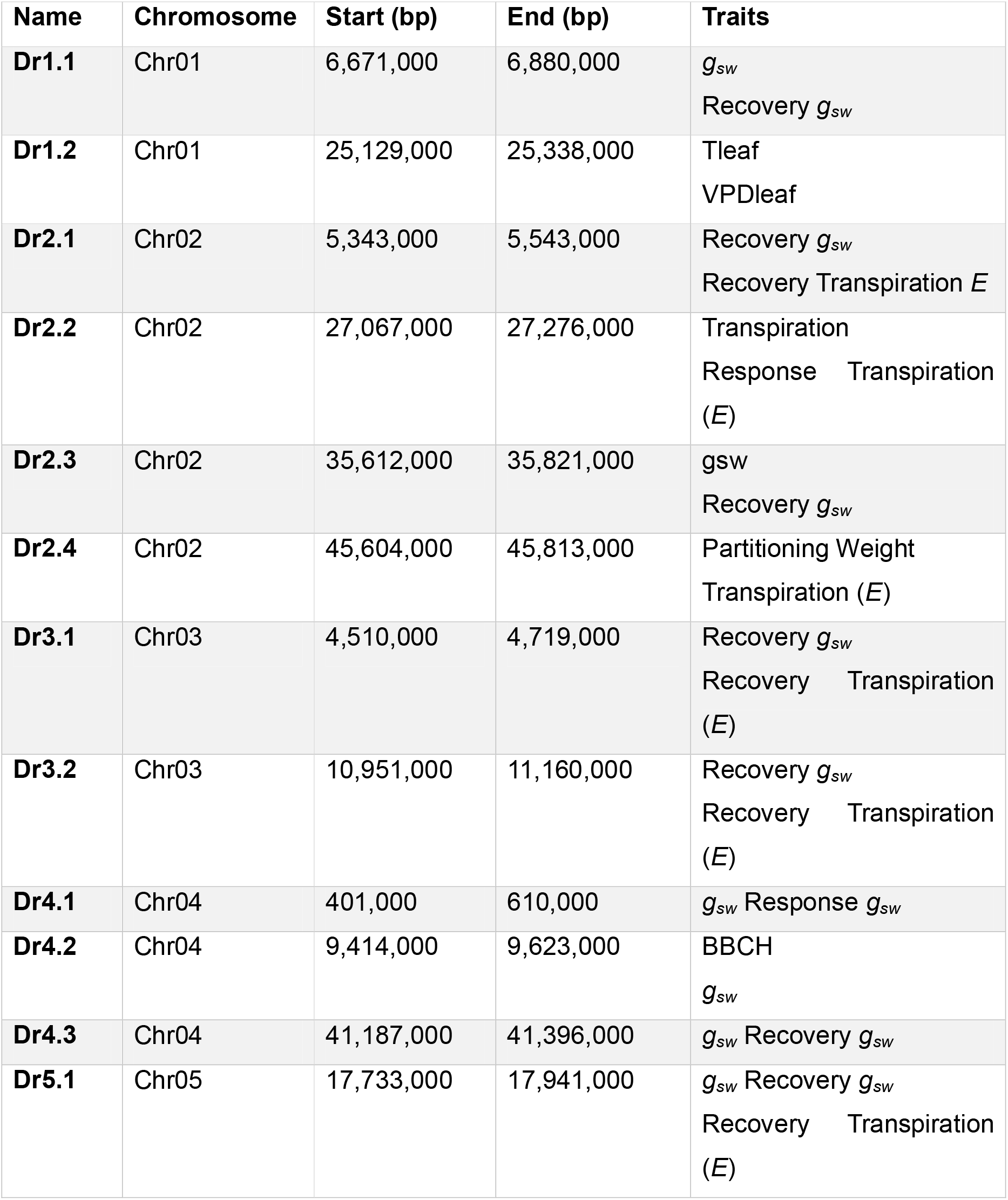
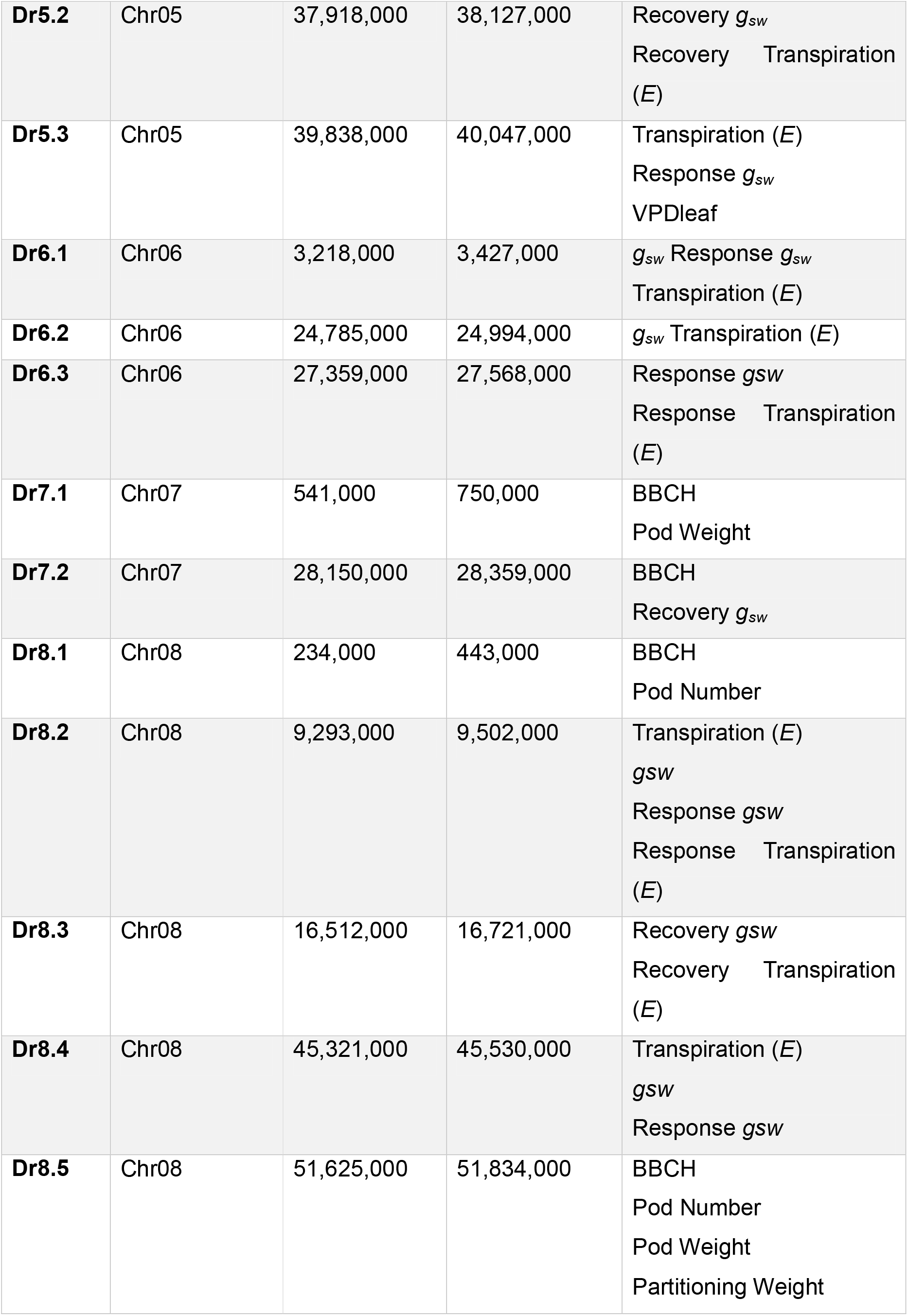
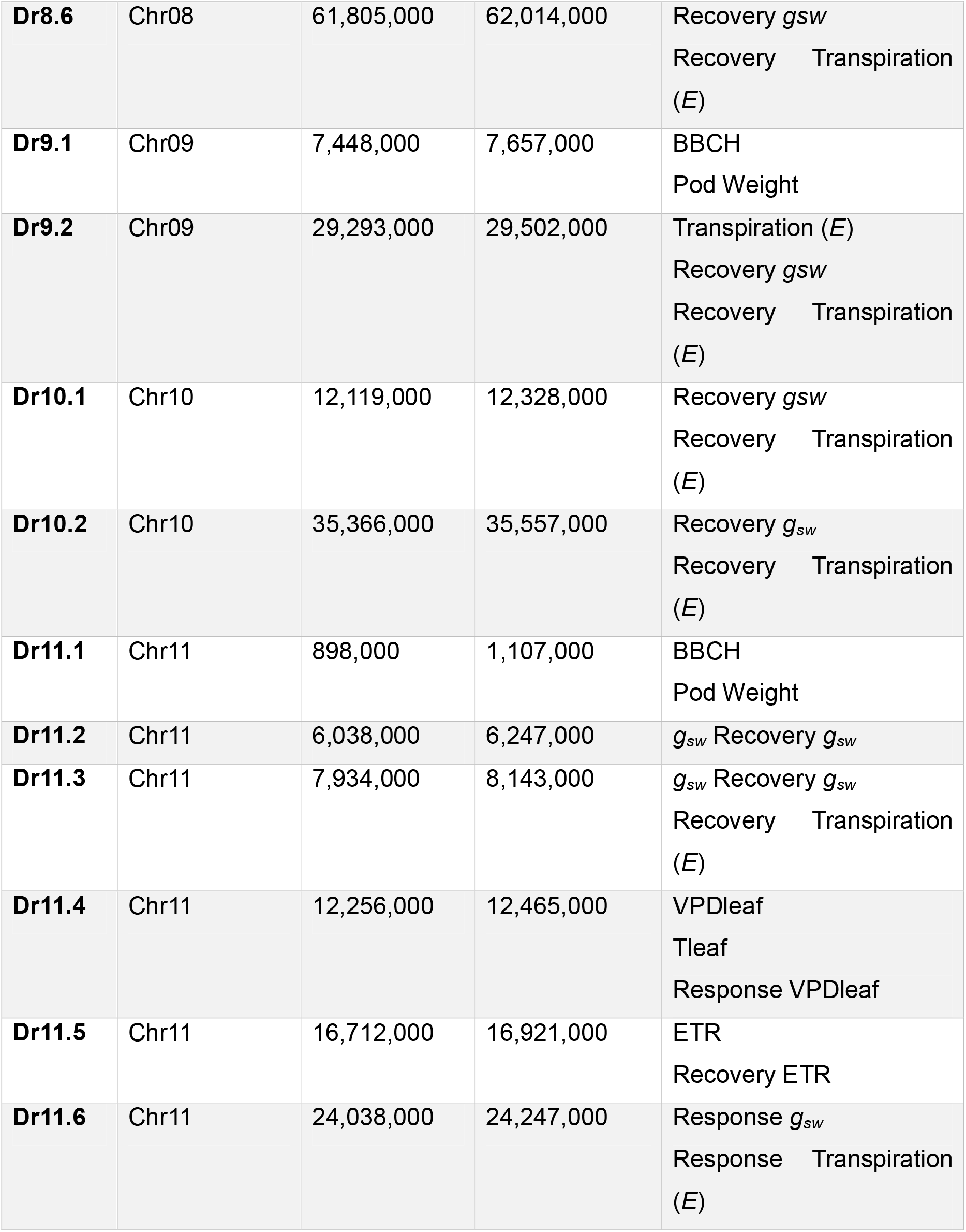
The 36 QTLs for *g_sw_* (porometer), *E* apparent (porometer), development (BBCH and yield), ETR (electron transport rate, fluorometer), VPDLeaf (leaf vapour pressure deficit), Tleaf (leaf temperature), water deficit ‘response’ (weeks 1 and 2) and ‘recovery’ from water deficit (weeks 4 and 5). Dr = “Drought-related QTL”.

| Name | Chromosome | Start (bp) | End (bp) | Traits |
| --- | --- | --- | --- | --- |
| **Dr1.1** | Chr01 | 6,671,000 | 6,880,000 | $g_{sw}$<br>Recovery $g_{sw}$ |
| **Dr1.2** | Chr01 | 25,129,000 | 25,338,000 | Tleaf<br>VPDleaf |
| **Dr2.1** | Chr02 | 5,343,000 | 5,543,000 | Recovery $g_{sw}$<br>Recovery Transpiration $E$ |
| **Dr2.2** | Chr02 | 27,067,000 | 27,276,000 | Transpiration<br>Response Transpiration<br>($E$) |
| **Dr2.3** | Chr02 | 35,612,000 | 35,821,000 | $g_{sw}$<br>Recovery $g_{sw}$ |
| **Dr2.4** | Chr02 | 45,604,000 | 45,813,000 | Partitioning Weight<br>Transpiration ($E$) |
| **Dr3.1** | Chr03 | 4,510,000 | 4,719,000 | Recovery $g_{sw}$<br>Recovery Transpiration<br>($E$) |
| **Dr3.2** | Chr03 | 10,951,000 | 11,160,000 | Recovery $g_{sw}$<br>Recovery Transpiration<br>($E$) |
| **Dr4.1** | Chr04 | 401,000 | 610,000 | $g_{sw}$ Response $g_{sw}$ |
| **Dr4.2** | Chr04 | 9,414,000 | 9,623,000 | BBCH<br>$g_{sw}$ |
| **Dr4.3** | Chr04 | 41,187,000 | 41,396,000 | $g_{sw}$ Recovery $g_{sw}$ |
| **Dr5.1** | Chr05 | 17,733,000 | 17,941,000 | $g_{sw}$ Recovery $g_{sw}$<br>Recovery Transpiration<br>($E$) |
| **Dr5.2** | Chr05 | 37,918,000 | 38,127,000 | Recovery $g_{sw}$<br>Recovery Transpiration<br>(E) |
| **Dr5.3** | Chr05 | 39,838,000 | 40,047,000 | Transpiration (E)<br>Response $g_{sw}$<br>VPDleaf |
| **Dr6.1** | Chr06 | 3,218,000 | 3,427,000 | $g_{sw}$ Response $g_{sw}$<br>Transpiration (E) |
| **Dr6.2** | Chr06 | 24,785,000 | 24,994,000 | $g_{sw}$ Transpiration (E) |
| **Dr6.3** | Chr06 | 27,359,000 | 27,568,000 | Response $g_{sw}$<br>Response Transpiration<br>(E) |
| **Dr7.1** | Chr07 | 541,000 | 750,000 | BBCH<br>Pod Weight |
| **Dr7.2** | Chr07 | 28,150,000 | 28,359,000 | BBCH<br>Recovery $g_{sw}$ |
| **Dr8.1** | Chr08 | 234,000 | 443,000 | BBCH<br>Pod Number |
| **Dr8.2** | Chr08 | 9,293,000 | 9,502,000 | Transpiration (E)<br>$g_{sw}$<br>Response $g_{sw}$<br>Response Transpiration<br>(E) |
| **Dr8.3** | Chr08 | 16,512,000 | 16,721,000 | Recovery $g_{sw}$<br>Recovery Transpiration<br>(E) |
| **Dr8.4** | Chr08 | 45,321,000 | 45,530,000 | Transpiration (E)<br>$g_{sw}$<br>Response $g_{sw}$ |
| **Dr8.5** | Chr08 | 51,625,000 | 51,834,000 | BBCH<br>Pod Number<br>Pod Weight<br>Partitioning Weight |
| **Dr8.6** | Chr08 | 61,805,000 | 62,014,000 | Recovery $g_{sw}$<br>Recovery Transpiration<br>($E$) |
| **Dr9.1** | Chr09 | 7,448,000 | 7,657,000 | BBCH<br>Pod Weight |
| **Dr9.2** | Chr09 | 29,293,000 | 29,502,000 | Transpiration ($E$)<br>Recovery $g_{sw}$<br>Recovery Transpiration<br>($E$) |
| **Dr10.1** | Chr10 | 12,119,000 | 12,328,000 | Recovery $g_{sw}$<br>Recovery Transpiration<br>($E$) |
| **Dr10.2** | Chr10 | 35,366,000 | 35,557,000 | Recovery $g_{sw}$<br>Recovery Transpiration<br>($E$) |
| **Dr11.1** | Chr11 | 898,000 | 1,107,000 | BBCH<br>Pod Weight |
| **Dr11.2** | Chr11 | 6,038,000 | 6,247,000 | $g_{sw}$ Recovery $g_{sw}$ |
| **Dr11.3** | Chr11 | 7,934,000 | 8,143,000 | $g_{sw}$ Recovery $g_{sw}$<br>Recovery Transpiration<br>($E$) |
| **Dr11.4** | Chr11 | 12,256,000 | 12,465,000 | VPDleaf<br>Tleaf<br>Response VPDleaf |
| **Dr11.5** | Chr11 | 16,712,000 | 16,921,000 | ETR<br>Recovery ETR |
| **Dr11.6** | Chr11 | 24,038,000 | 24,247,000 | Response $g_{sw}$<br>Response Transpiration<br>($E$) |

For convenience, the QTLs were grouped in categories based on the capture method or complex trait associated: water vapour measured by porometer (averaged, differences between weeks, total, and scaled; figure 6A), transpiration by porometer (similar transformations; figure 6B), BBCH and harvest weights and traits (figure 6C), ETR by fluorometer and leaf T and VPD by porometer (figure 6D), “early response” evaluated in weeks 1 and 2 (figure 6E) and “recovery response” evaluated in weeks 4 and 5 (figure 6F).

Across the thirty-six QTLs selected across all traits (table 4), twelve QTLs were found associated with stomatal conductance (*g_sw_*) in chromosomes Pv01, Pv02, Pv04, Pv05, Pv06, Pv08 and Pv11 (figure 6A), and eight QTLs were associated to transpiration (*E*) in chromosomes Pv02, Pv05, Pv06, Pv08 and Pv09 (figure 6B). Eight QTLs were found associated to developmental traits (BBCH and harvesting) in chromosomes Pv02, Pv04, Pv07, Pv08, Pv09 and Pv11 (figure 6C), and ETR, VPDleaf and Tleaf were associated with three QTLs on chromosomes Pv01 and Pv11. VPDleaf and Tleaf share the same two QTLs and are significantly correlated (figure 6D). Eight QTLs were related to “early response” in chromosomes Pv02, Pv04, Pv05, Pv06, Pv08 and Pv11 (figure 6E), and seventeen QTLs were identified related to “recovery response” in chromosomes Pv01, Pv02, Pv03, Pv04, Pv05, Pv07, Pv08, Pv09, Pv10 and Pv11 (figure 6F).

Within these 36 QTLs there were 465 genes (Supplementary table S3). After prioritising genes within the QTLs annotated with GO terms related to drought or water deficit stress, we identified thirteen promising candidate genes: Phvul.001G058600, Phvul.002G288700, Phvul.004G007100, Phvul.004G122000, Phvul.005G138000, Phvul.006G005100, Phvul.006G170600, Phvul.006G171100, Phvul.007G167200, Phvul.008G092800, Phvul.008G275300, Phvul.009G192900 and Phvul.011G013900. Among them, four genes (Phvul. 001G058600, Phvul. 004G122000, Phvul. 005G138000, Phvul. 011G013900) contained ‘high impact’ non-synonymous (frameshift) mutations.

## Discussion

### Indicators of water-deficit stress

Soil moisture sensors provided a quantitative measure of the water deficit stress experienced by the plants. A soil matric potential lower than-800 kPa and a water content below 0.1 m^3^/m^3^, as recorded in our experiment (figure 1), indicate water deficit and are sufficient to induce a drought stress response (Ganesan et al. 2024; Gebregiorgis and Savage 2006; Lui and Mihara 2024; Medynska-Juraszek et al. 2021; Yang et al. 2024). Consequently, the limited water supply to the accessions in the evaluation experiment failed to meet the requirements for normal growth and development (Dramadri et al. 2019; Wortmann et al. 1998).

Overall, there was no significant statistical difference in soil water content among determinate bush, indeterminate bush, or indeterminate climbing plants under water deficit conditions. The only exception was soil water content and Bulk EC at week 2 (supplementary figure S3), indicating that differences in vegetative growth might have caused indeterminate accessions to access more soil water earlier after the treatment was applied (Campos *et al*. 2021).

Because the UK climate is mild, water deficit was observed in our experiment without strong heat stress, as soil and air temperatures remained within normal ranges. This improves interpretability for traits such as stomatal conductance, leaf temperature and developmental progression. However, combined stresses are likely more common where common bean is a main crop (Sato *et al*. 2024). The trial used pots to better control water application. While field trials under rainout shelters arguably more closely resemble real-world conditions, they are logistically complex and hard to standardize due to variable rainfall, soil, and plot differences (Beebe *et al*. 2013; Langstroff *et al*. 2022). Pot systems can offer better control over substrate, plant density, irrigation, and stress timing, enabling repeated measurements at the individual-plant level. This is especially valuable for studies on response dynamics rather than final yield. However, pot-based methods also influence root structure and root plasticity is linked to drought response. Some accessions may develop more extensive root systems, and indeterminate types, which are usually larger, may be more affected in open soils (Velho *et al*. 2018; Cerutti *et al*. 2023).

One of our objectives was to interpret quantitative plant responses to decreasing water availability using leaf porometry and fluorometry. The steady state fluorescence (Fs) and maximum fluorescence yield (Fm’) declined from week 1 to week 4, consistent with altered PSII photochemistry under stress (Guidi *et al*. 2019; Ramírez-Estrada *et al*. 2023; Jat *et al*. 2024). Additionally, higher leaf temperatures (Tleaf) during drought show a negative correlation with other leaf traits, leading to reduced stomatal conductance (*g_sw_*) and lower photosynthetic efficiency, such as decreased ETR and Fm’. This further supports the use of leaf temperature as an indicator of stomatal conductance under drought conditions in common bean, a method utilised in canopy-level infrared thermal imaging (Yu *et al*. 2015; Driever *et al*. 2023). The plant’s phenological development also negatively correlates with stomatal and photosynthetic traits (Cavalcante *et al*. 2020). This is likely because the BBCH scale is confounded by pod development, whereas drought-induced maturation hampers stomatal conductance and transpiration (*g_sw_* and *E*_apparent, respectively) (Müllers *et al*. 2022; Jahan *et al*. 2023; Urban and Urban 2024).

### Response strategies

The panel includes accessions that are susceptible to drought, as they exhibit low pod and foliar weights after four weeks of water deficit. Common beans are known to suffer significant yield losses under drought stress, underscoring the importance of identifying those with distinct drought-tolerance strategies (Rosales *et al*. 2012; Labastida *et al*. 2023). Yet, we identified multiple drought tolerance strategies within the diversity panel, categorised as ‘stay-green’, ‘saver’, ‘spender’, ‘prioritised yield’, ‘drought susceptible’, ‘nominal growth’, and ‘no classification’. These strategies were found in both gene pools (with one exception due to limited sampling), likely because of local adaptation to the specific agro-ecogeographic environment.

These different strategies have been shaped by adaptation to local agro-ecogeographic conditions and farming methods (Blum 2015; Polania *et al*. 2016). All strategies can consequently be beneficial for breeding programmes to tailor cultivars to the prevalent type of drought in a region. Our work found that developmental information was critical to distinguish strategies, and when complemented with porometer and fluorometer data (mainly Tleaf, *g_sw_*, and *E*_apparent) can indicate the level of water deficit stress the plants are experiencing. By contrast, these data alone were unable to differentiate among drought tolerance strategies; e.g., accessions with very high photosynthetic efficiencies (top 10%) encompass a variety of response strategies, including a few susceptible accessions.

‘Savers’ (isohydric) reduce gas exchange and photosynthesis under drought conditions by closing their stomata earlier (Polania *et al*. 2016). These accessions halt development or abort pod formation to wait for better conditions. In our panel, this was shown by the BBCH stage not advancing (pausing) during water deficit, therefore not reaching pod development, and sometimes resuming during the ‘recovery’ phase. Without additional watering, savers typically yield fewer seeds (Hamabwe *et al*. 2024), as shown in our accessions. This strategy is better suited to prolonged mild droughts, in which water returns, allowing the plant to produce seeds rather than just vegetative biomass (Bandurska 2022; Nesporová *et al*. 2024). A large portion of the panel were ‘savers’, indicating that this strategy may be better adapted to the accessions’ and/or species’ original ecogeographic environment (Arregocés *et al*. 2025).

‘Spenders’ (anisohydric) maintain gas exchange and photosynthesis, accelerating growth under stress conditions to avoid the onset of worsening conditions. In our panel, this was shown by continued development (increasing BBCH) alongside high transpiration and photosynthesis rates. The spending strategy is most effective for producing yields during long-term drought stress (Delfin *et al*. 2021; Nesporová *et al*. 2024). If the accessions can complete their life cycle before conditions deteriorate, this approach is ideal for longer-term drought stress when rainfall may not return, similar to drought escape (Shavrukov *et al*. 2017). However, if the lifecycle is not completed, this strategy may lead to total crop failure. A subset of these includes those which ‘prioritised yield’, maintaining fertilisation and grain development under drought stress (Polania *et al*. 2016). In our panel, these had seed yields in the top 10% under treatment, indicating they could be considered tolerant to drought (Hamabwe *et al*. 2024).

“Stay-green” accessions sustain photosynthesis, while delaying senescence and chlorophyll degradation, under drought stress, enabling rapid resumption of development when water is available again (Thomas and Ougham 2014). In our trial, they exhibited very high foliar weight but very low or zero pod weight at harvest. This strategy is most effective for shorter droughts when water returns quickly. Further research is needed to evaluate the yield of stay-green accessions under drought stress and during recovery compared with controls. In other crops, the ‘stay-green’ strategy has been associated with the highest yields under drought conditions (Kamal *et al*. 2019; Padilla-Chacón *et al*. 2019). In our panel, all stay-green accessions were Andean; however, they have been previously identified in other common bean populations (Schmit *et al*. 2019; Labastida *et al*. 2023).

Notably, there was also a correlation between seed colour and water deficit strategies. This has been reported previously in common bean (Hussaini *et al*. 2021), likely because in-country market preferences and climate act as confounding factors, similar to the high correlation observed among seed colour, seed weight, and seed size with gene pools (Giordani *et al*. 2022).

Additional traits could enhance the above-ground traits analysed, which have been linked to drought response estimation, including the stay-green trait, rates of photosynthesis, transpiration, and reproductive output (Sofi *et al*. 2021). Root traits are strongly associated with response strategies (Polania *et al*. 2017). Roots typically react first to drought stress in plants; however, because belowground sampling and observations are challenging, foliar organs are often studied instead (Irshad *et al*. 2024). A detailed examination of epidermal traits, such as stomatal density (Egesa *et al*. 2024) and size (Polania *et al*. 2022), may also provide valuable insights.

### Recovery response

Precipitation (rainfall) increased after the measurements in week 4 (Supplementary Figure S2), leading to higher soil water content, as plants were not sheltered from rainfall. In week 5, soil water content (m^3^/m^3^) was equivalent to that observed in week 3. This precipitation was insufficient to fully offset the water previously lost, as supported by the continued decline in soil metrics at week 5 (Figure 1, Supplementary Figure S3), but it was sufficient to induce a “recovery response” in the treated plants, both photosynthetically and developmentally, as supported by the significant differences between weeks 4 and 5 in some traits (Supplementary Figure S8). One of the traits that did not significantly change was plant development (scored using BBCH, Figure 4), likely because plants that had not flowered did not have enough time to accelerate their development, for example towards flowering, and plants that had flowered during the water deficit had already begun producing pods or were maturing. How crop plants recover photosynthesis and growth after water deficit affects their ability to continue normal development and, consequently, yields, as drought is often intermittent or can be partially alleviated through irrigation and other water management practices (Wang *et al*. 2019; Delfin *et al*. 2021). A relevant research direction would be to evaluate ‘stress memory’, the “imprinting” of past stress events in future stress responses (Jacques *et al*. 2021).

### Candidate genes

By analysing this diversity panel, 36 QTLs were identified as being associated with responses and recovery to water deficit (Table 4). Among these QTLs, there were 465 genes (supplementary table S3). Gene ontology terms were subsequently filtered to select genes associated with tolerance to water stress.

One candidate gene, Phvul.001G058600 (QTL Dr1.1), encodes a protein involved in chloroplastic chaperone activity of the BC1 complex (CABC1) (Goodstein *et al*. 2012), and may modulate chlorophyll degradation in response to oxidative stress, including salt stress (Huala *et al*. 2001; Borkiewicz *et al*. 2020; Qin *et al*. 2020). Other drought-stress-associated genes were identified: Phvul.005G138000 (Dr5.2) is orthologous to Arabidopsis *NPF2.6* (AT3G45660), a member of the NAXT NPF subfamily which is upregulated under drought and involved in xylem transport in response to osmotic stress (Huala *et al*. 2001; Li *et al*. 2010, 2016; Morales de Los Ríos *et al*. 2026); Phvul.011G013900 (Dr11.1) is likely to encode a protein orthologous to Arabidopsis calcium transporting ATPase11 (AT3G57330.1) (Huala *et al*. 2001; Goodstein *et al*. 2012), associated with drought and salt tolerance, possibly via stomatal control (He *et al*. 2024; Su *et al*. 2024).

Transcription factor-encoding genes in the QTLs include Phvul.004G122000 (Dr4.3), which may encode *DREB1*, a dehydration response element binding protein (Goodstein *et al*. 2012). DREBs are water-deficit response transcription factors common across the plant kingdom (Han *et al*. 2022), such as rice (Wang *et al*. 2022), wheat (Mei *et al*. 2022), tomato (Tao *et al*. 2022). Another DREB gene, Phvul.008G092800 (orthologous to *DREB2*) was also found within the QTLs (*Dr8.2*). Both *DREB1* and *DREB2* are involved in responses to abiotic stresses, including heat, drought and salt (Akhtar *et al*. 2012; Guttikonda *et al*. 2014; Akbudak *et al*. 2018). Other TFs include SHINE orthologue Phvul.010G092300 (Dr10.2), encoding an APETALA2/ Ethylene Responsive Factor (AP2/ERF) controlling waxy cuticle and stomatal development (Girón-Ramírez *et al*. 2021; Khoudi 2023), MYBs (Phvul.004G121500, Phvul.009G192600, Phvul.011G084500, Dr4.3, Dr9.2, and Dr11.3, respectively), and Phvul.008G275300 (Dr8.6), an orthologue of *WRKY20*, known to modulate guard cell ABA and reactive oxygen species (ROS) signalling as well as cuticular wax biosynthesis in response to drought (Rushton *et al*. 2012; Luo *et al*. 2013; Li *et al*. 2023). ERFs, MYBs, WRKYs, and protein kinases are gene families that respond to drought stress and to drought tolerance mechanisms by interacting with downstream genes in signalling pathways (Umezawa *et al*. 2006; Wu *et al*. 2024).

Genes related with the phytohormones auxin (Phvul.005G172900, Phvul.005G173000, Phvul.006G142200, Phvul.006G142300) (Liu *et al*. 2019; Wang *et al*. 2021; Yin *et al*. 2023) and gibberellic acid (Phvul.002G193300, Phvul.005G173500, Phvul.007G167600) were also identified (Chu *et al*. 2022). Phvul.007G167600 is an orthologue of *GIBBERELLIN-INSENSITIVE DWARF 1* (*GID1*), a partially ABA-dependent GA receptor that regulates stomatal development and ABA biosynthesis under drought stress in rice (Du *et al*. 2015). Other ABA-associated genes identified in these pathways include those encoding pentatricopeptide repeat proteins (PPRs) (Phvul.001G059500, Phvul.003G071600, Phvul.008G277600, Phvul.009G034500, Phvul.011G068200, Phvul.011G069000) including a DYW subgroup PPR gene, (Phvul.011G107600) linked to soybean drought responses (Su *et al*. 2019). Similarly, orthologues to protein kinase-encoding genes involved in drought and ROS signalling were identified (Phvul.002G289000, Phvul.005G173100, Phvul.008G093200, and Phvul.008G124300) (Xu *et al*. 2018; Liu *et al*. 2020, 2024; Wu *et al*. 2024; Cai *et al*. 2025). Enhanced photosynthetic efficiency under drought may arise from superior responses to oxidative stress (ROS). Together with transcription factors, crosstalk among auxin, gibberellic acid, and ABA regulates an array of signalling components in downstream pathways critical to water-deficit responses (Salehin *et al*. 2019; Aslam *et al*. 2022; Verma *et al*. 2022; Labastida *et al*. 2023; da Silva *et al*. 2024).

The candidate genes identified in this diversity panel appear to be important for drought-stress resilience. However, additional functional tests are needed to verify whether these potential genes play similar roles in common bean as their orthologues do in other crops such as soybean and Arabidopsis. Since most QTLs were identified using porometry and fluorometry measurements, integrating these metrics into future research will be crucial for detecting and comparing variation in stomatal and photosynthetic drought responses, alongside developmental data.

## Conclusions

This study demonstrates that common bean exhibits substantial diversity in integrated above-ground responses to water deficit, spanning developmental progression, biomass allocation, stomatal and photosynthetic performance, and short-term recovery after stress relief. The principal contribution of this work is the use of a panel-scale, integrative phenotyping framework to resolve partially overlapping response profiles under a defined pre-flowering water-deficit scenario, rather than introducing fundamentally new drought-response concepts. Developmental data were especially important for interpreting physiological measurements and for distinguishing accessions that differed in stress progression and recovery. By linking these response profiles and their component traits to population structure, gene flow and GWAS signals, this study provides candidate loci, trait relationships and a practical framework for future validation across environments and for breeding strategies aimed at improving adaptation to variable water availability under climate change.

## Data availability statement

Raw reads are deposited in the SRA under accession PRJEB81566 in our previous publication (Denning-James *et al*. 2025). The scripts used in this study are publicly available on GitHub (https://github.com/DeVegaGroup/KDJ-CBeans/).

## Acknowledgments

All the authors approved this manuscript. We gratefully acknowledge the essential support from the technical teams at Niab Park Farm at Niab (Cambridge, UK) and the Transformative Genomics group at the Earlham Institute (Norwich, UK), for their extensive assistance with phenotypic and genotypic data collection, respectively. We also thank the Earlham Institute’s research office, BDE and support teams, as well as the Norwich Bioscience Institute’s research computing and contracts teams, and horticultural services at the John Innes Centre. We thank CIAT’s Genebank and IPK’s Genebank for generously providing germplasm.

---

**Supplementary Table S1:** Phenotypic traits recorded at multiple time points for a diversity panel of common bean (*Phaseolus vulgaris* L.) accessions.

**Supplementary Table S2:** Phenotypic scores used for GWAS

**Supplementary Table S3:** Annotation of the genes within the QTLs.

