## supplementary figure s1 for "Developmental and physiological profiles define drought response diversity and genomic associations in common bean"

**Supplementary Figure S1:** A water-deficit experiment was performed outdoors at the NIAB research station (Histon, UK; 52.246, 0.098) from June to September 2023.

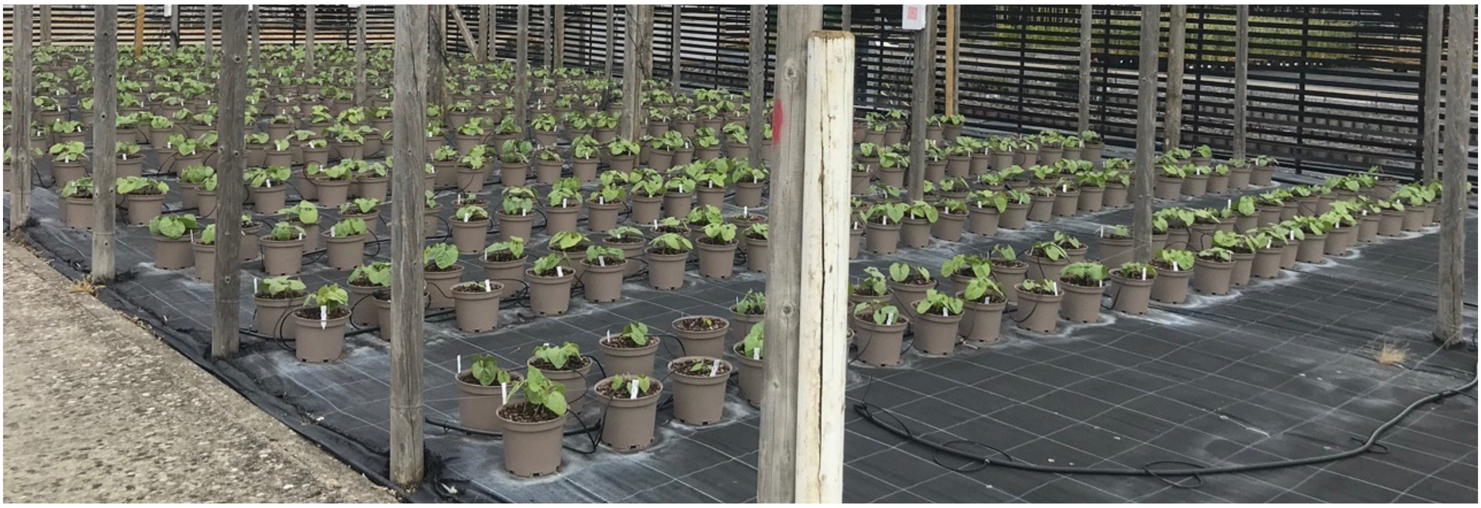
