## supplementary figure s2 for "Developmental and physiological profiles define drought response diversity and genomic associations in common bean"

**Supplementary Figure S2: A water-deficit experiment was performed outdoors at the NIAB research station (Histon, UK; 52.246, 0.098) from June to September 2023.** Irrigation was stopped on day 38 after sowing, subsequently subjected to water-deficit for five weeks (weeks 1 to 5). Throughout the experiment, the average day length was 15.4 hours, and the ambient temperature averaged 17.6 °C. The mean precipitation between the time when irrigation stopped (day 38) and harvest (day 79) was 1.73 mm/day.

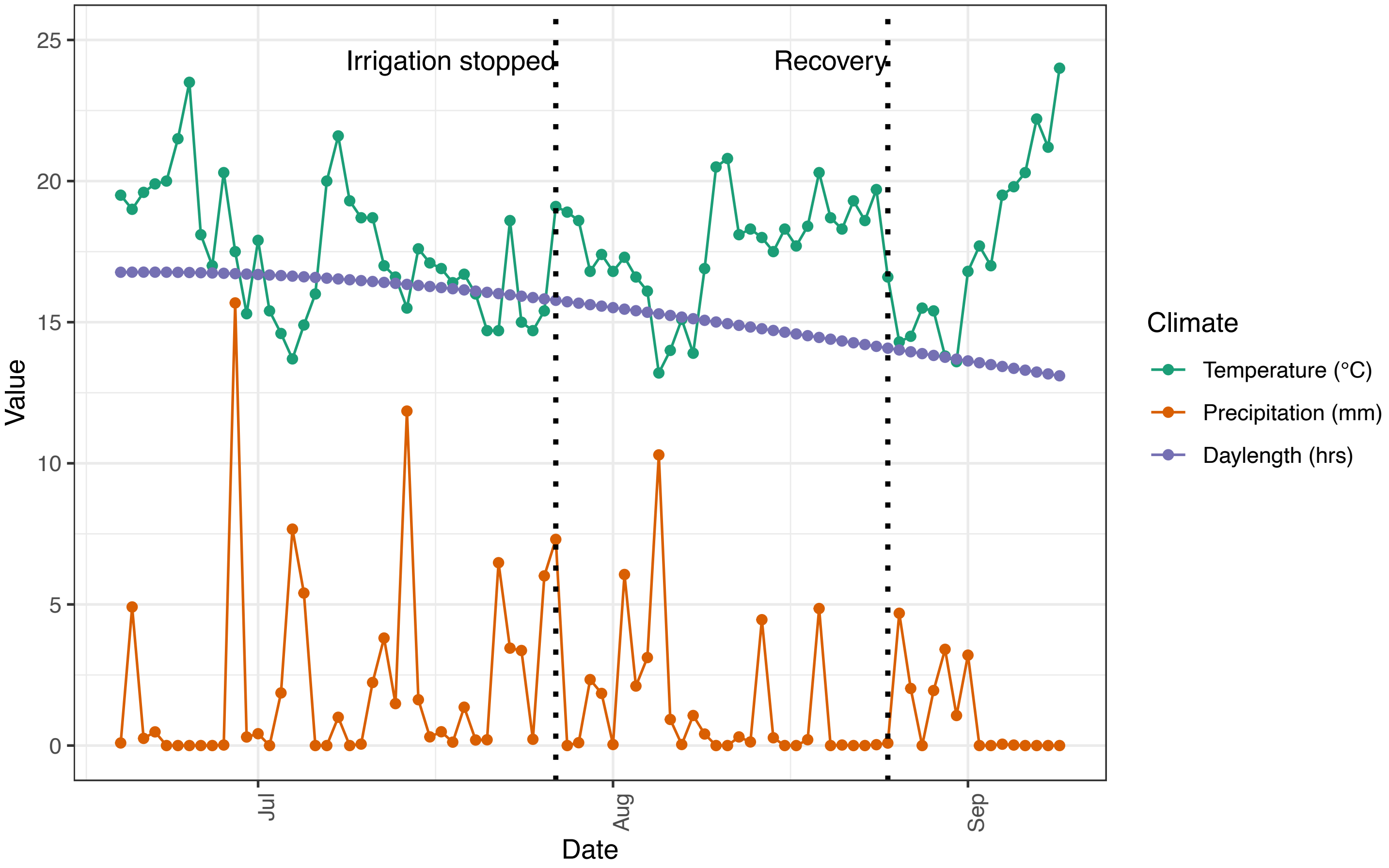
