## supplementary figure s3 for "Developmental and physiological profiles define drought response diversity and genomic associations in common bean"

**Supplementary figure S3: Measured parameters collected from soil moisture sensors and grouped by growth habit and stress condition (control or treated).** (A) Soil Matric Potential (kPa), (B) soil temperature (°C), (C) the bulk electrical conductivity (EC, mS/cm) and (D) for the water content ($m^3/m^3$). Panels correspond to collecting date, from irrigation (week 0) to harvesting (week 5). Statistical analyses identified significant interactions between stress x growth habit over time. Groups were labelled by a letter and superscript number, where groups with different letters differed significantly at α = 0.05, and the superscript number corresponded to the week of collection.

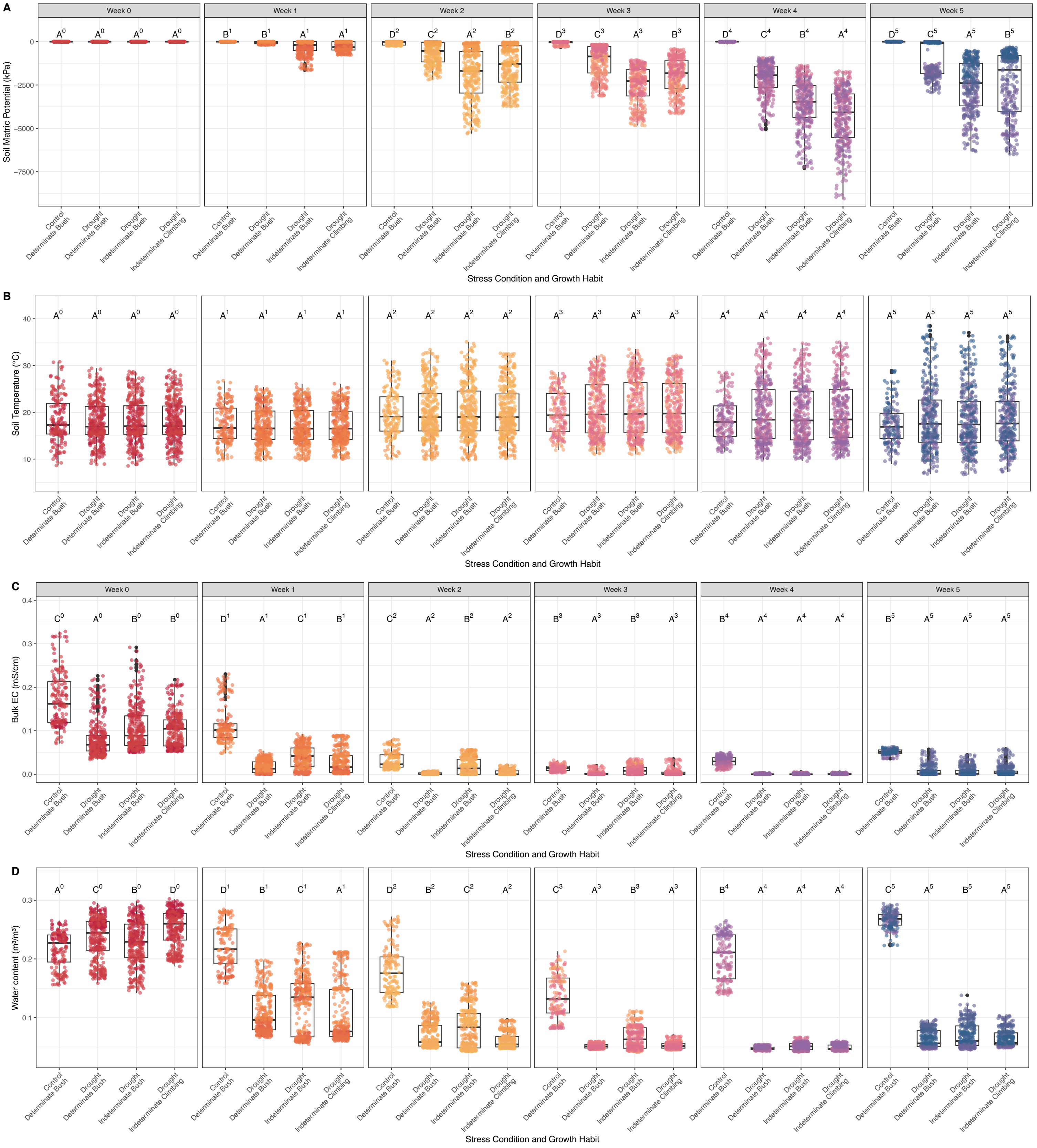
