## supplementary figure s4 for "Developmental and physiological profiles define drought response diversity and genomic associations in common bean"

**Supplementary figure S4: Measured scores collected from the porometer and fluorometer for each accession and grouped by growth habit.** Panels correspond to collecting date (week 1 to week 4). (A) Fm' (maximum fluorescence yield), (B) Fs (steady-state chlorophyll fluorescence), (C) Tleaf (leaf temperature, °C), (D) GSW ($g_{sw}$, stomatal conductance, mol m$^{-2}$s$^{-1}$), (E) E apparent ($E$, transpiration, mmol m$^{-2}$s$^{-1}$) and (F) PhiPS2 (ΦPSII, quantum yield of fluorescence). Groups sharing the same letter are not significantly different at α = 0.05, whereas groups with different letters differed significantly. The superscript numbers correspond to the week.

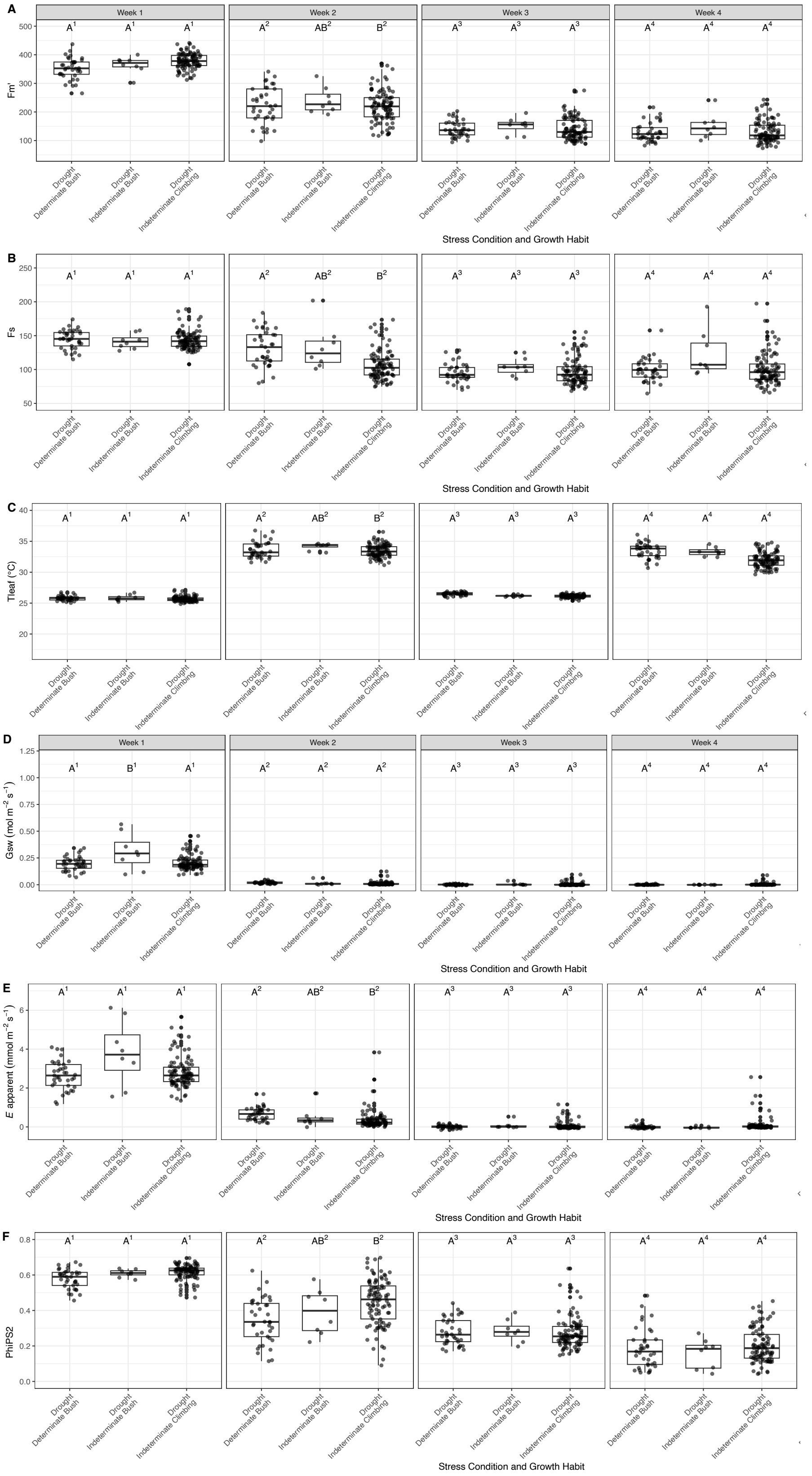
