## Supplementary figures and images for "Developmental and physiological profiles define drought response diversity and genomic associations in common bean"

### supplementary figure s5

**Supplementary figure S5: Stomatal conductance vapour ($g_{sw}$) (mol/m²/s)**

A

**Week 0**

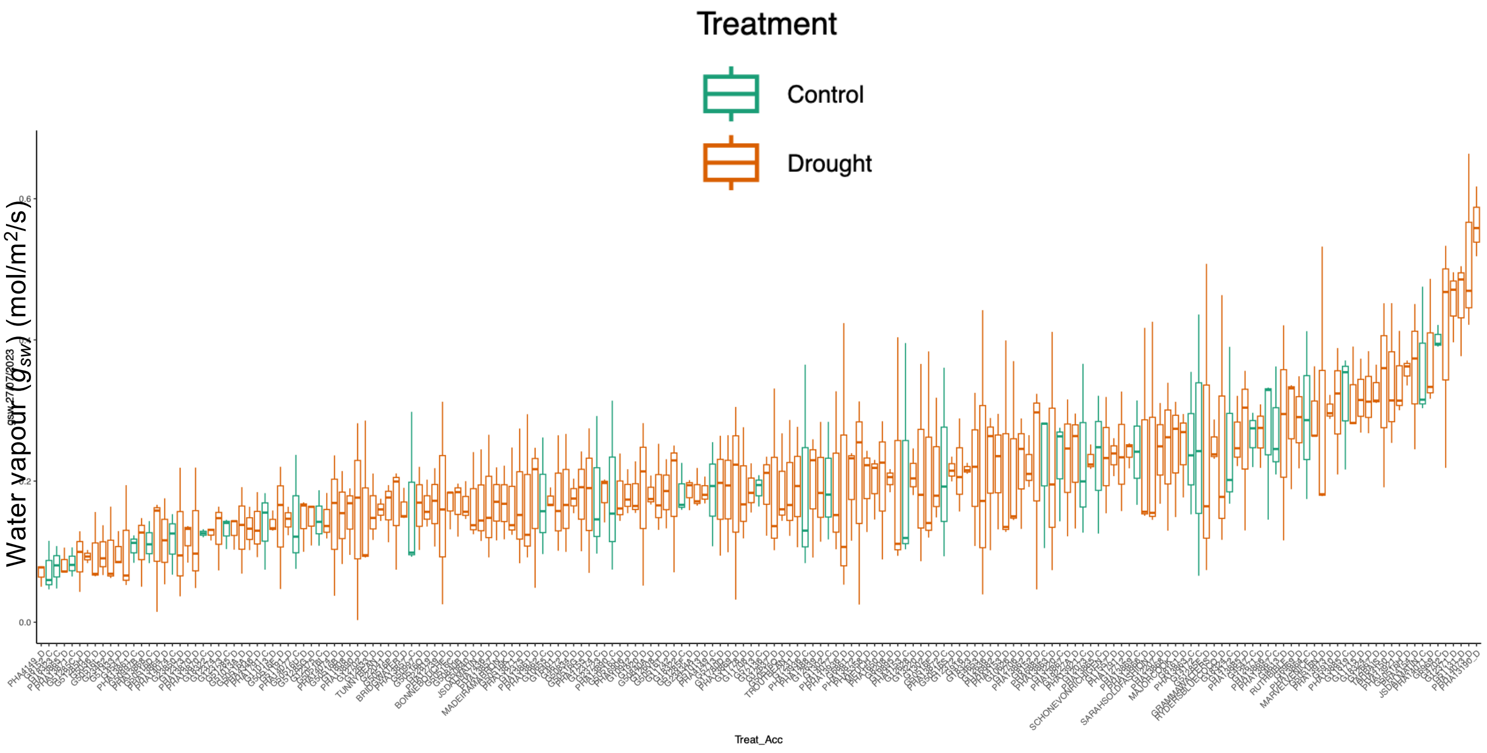


B

**Week 2**

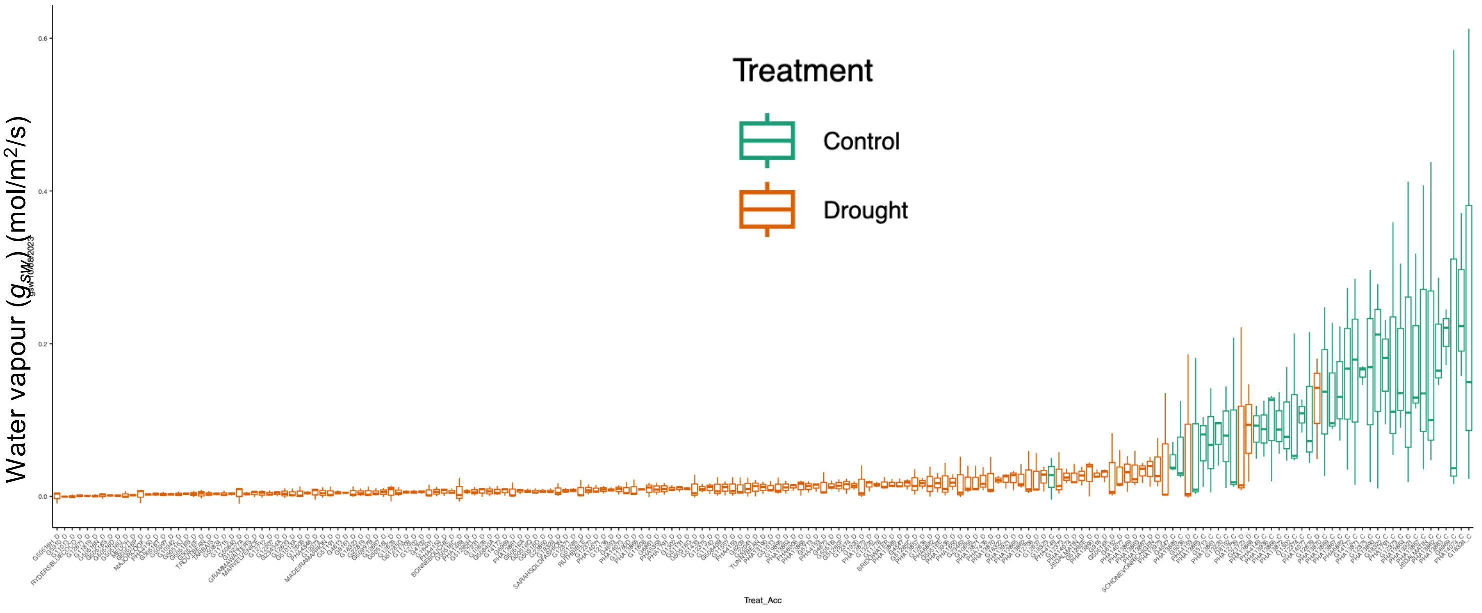

### supplementary figure s6

#### Supplementary figure S6: normalised leaf T (Tleaf) ℃

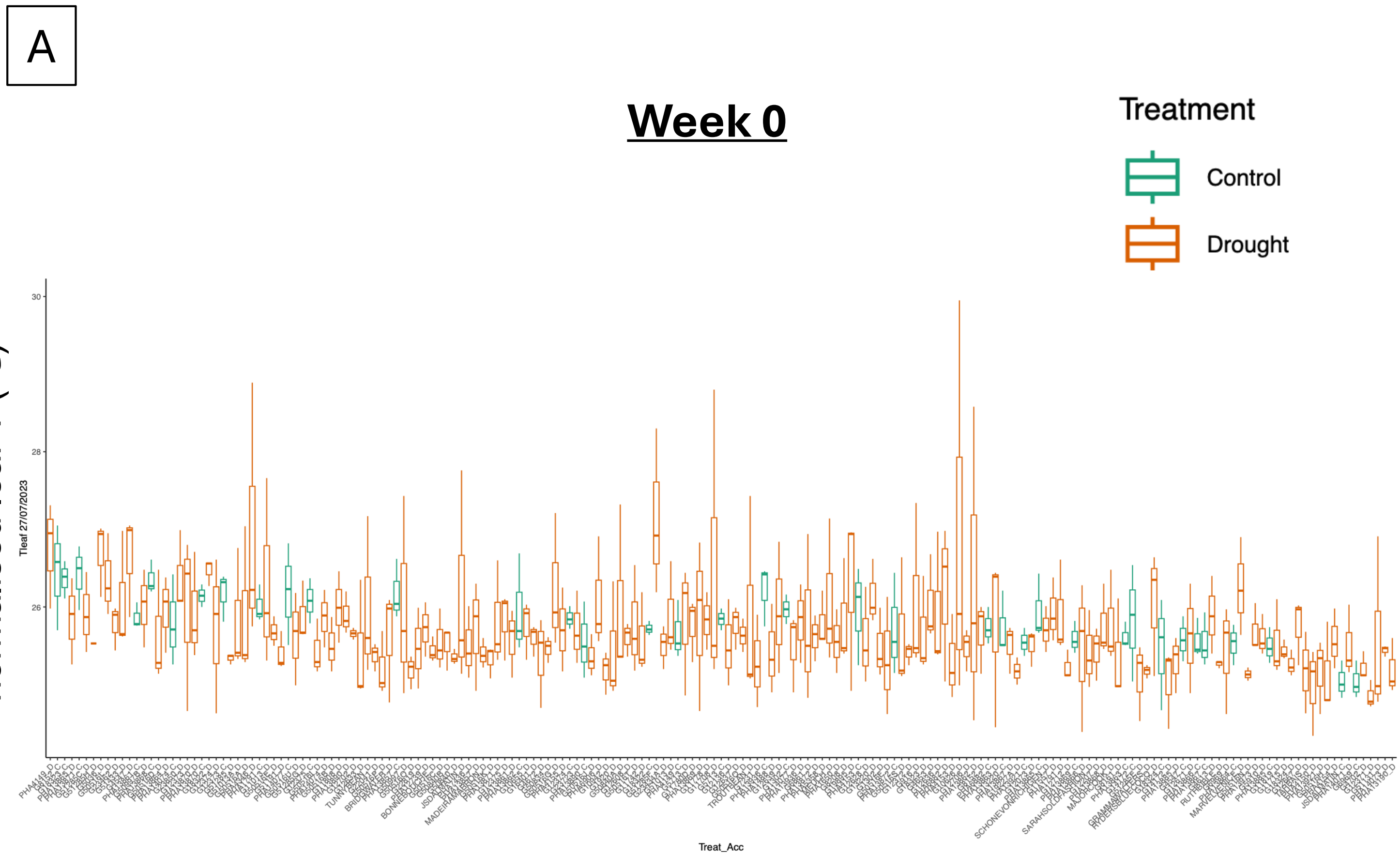


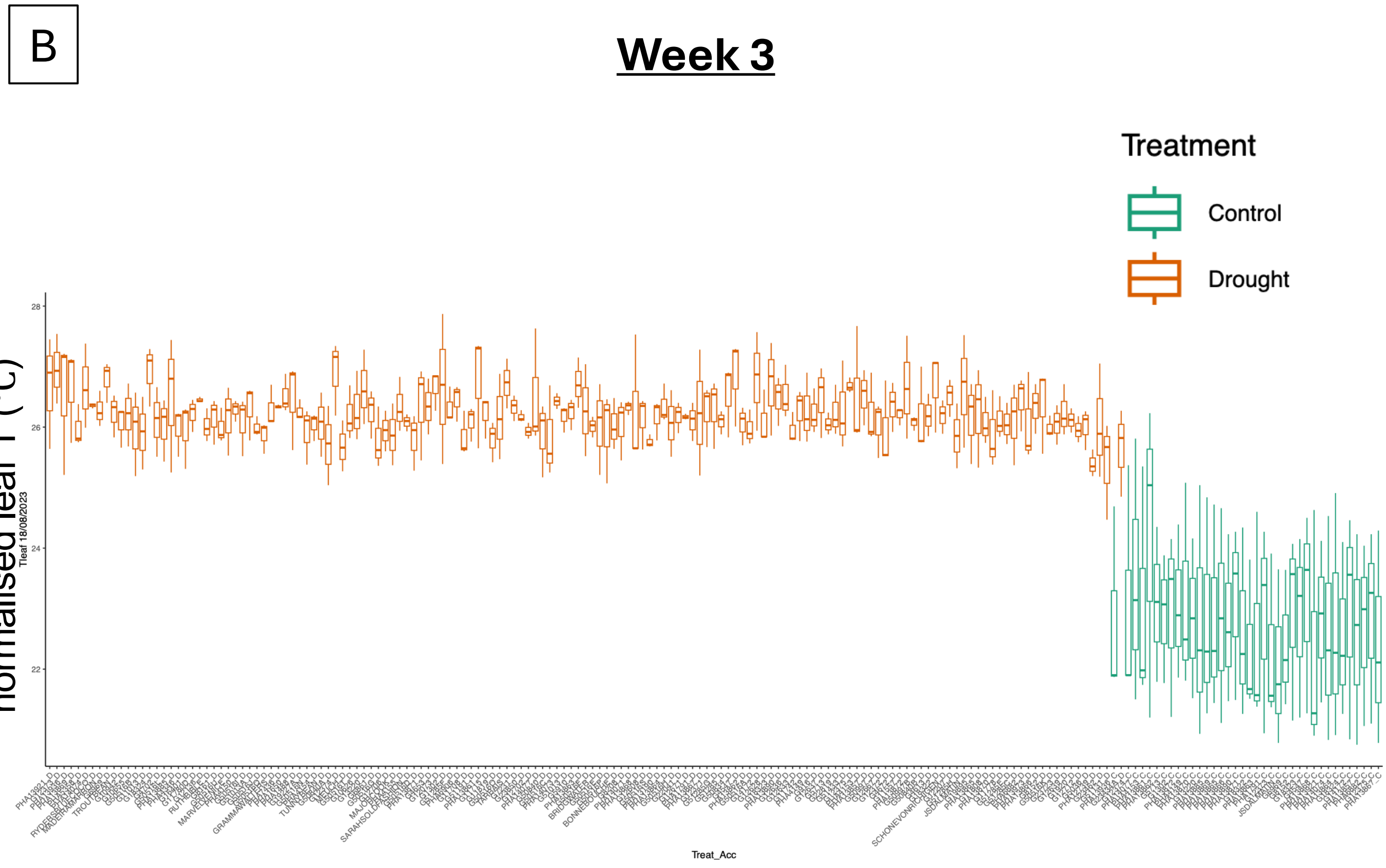
