## supplementary figure s7 for "Developmental and physiological profiles define drought response diversity and genomic associations in common bean"

### Supplementary figure S7: Comparison of trait values scored per accession across growth habit and condition

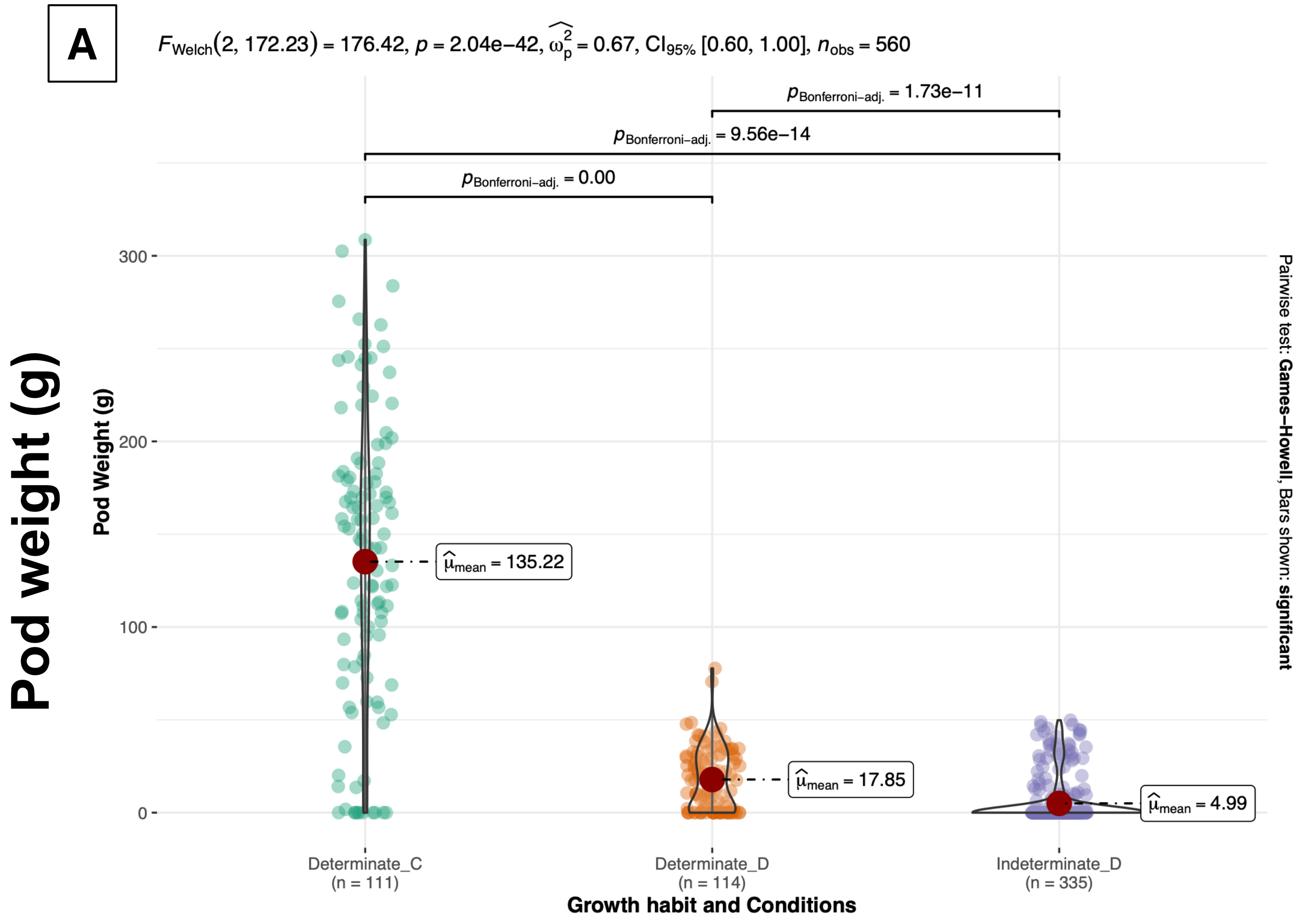


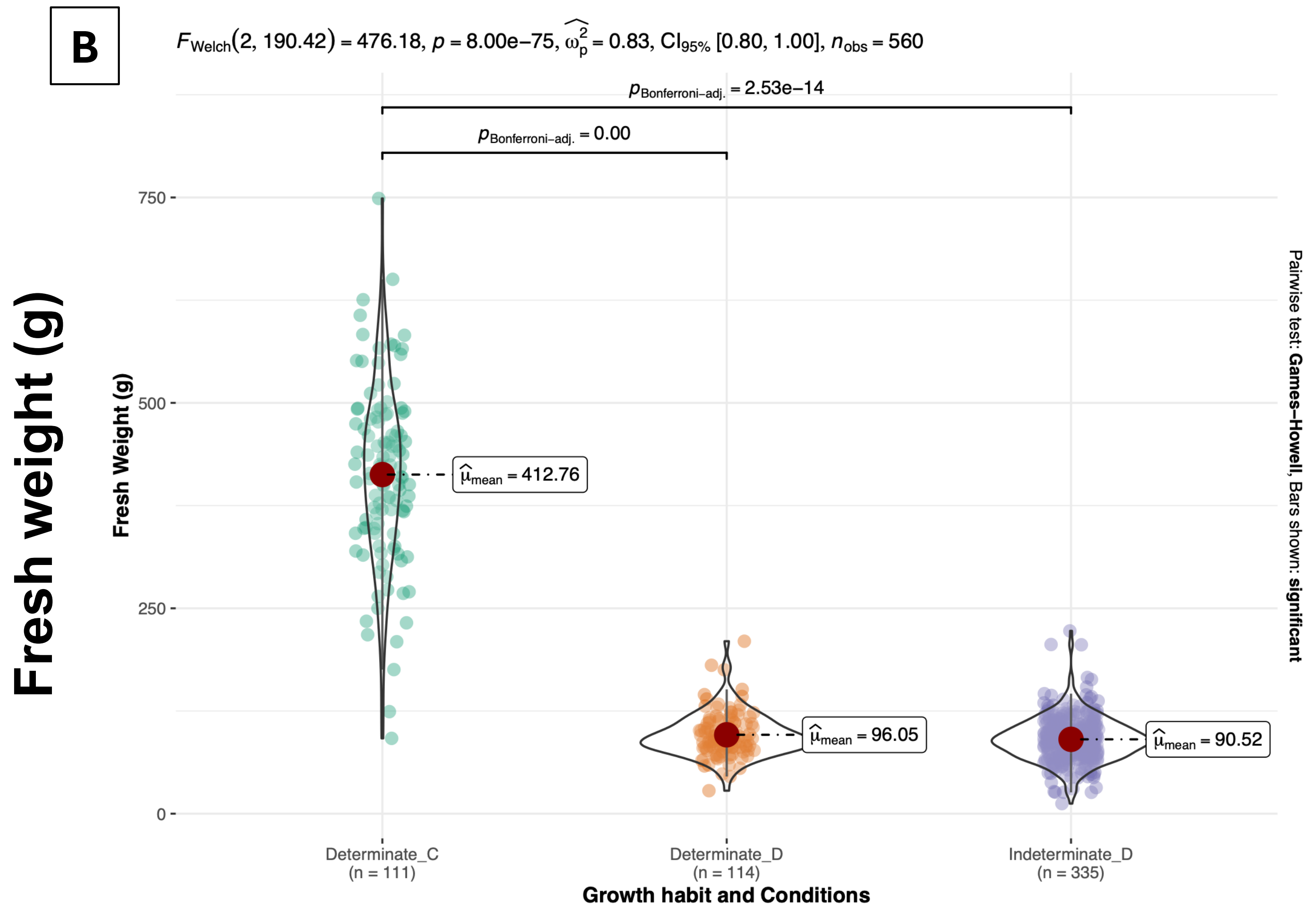

C

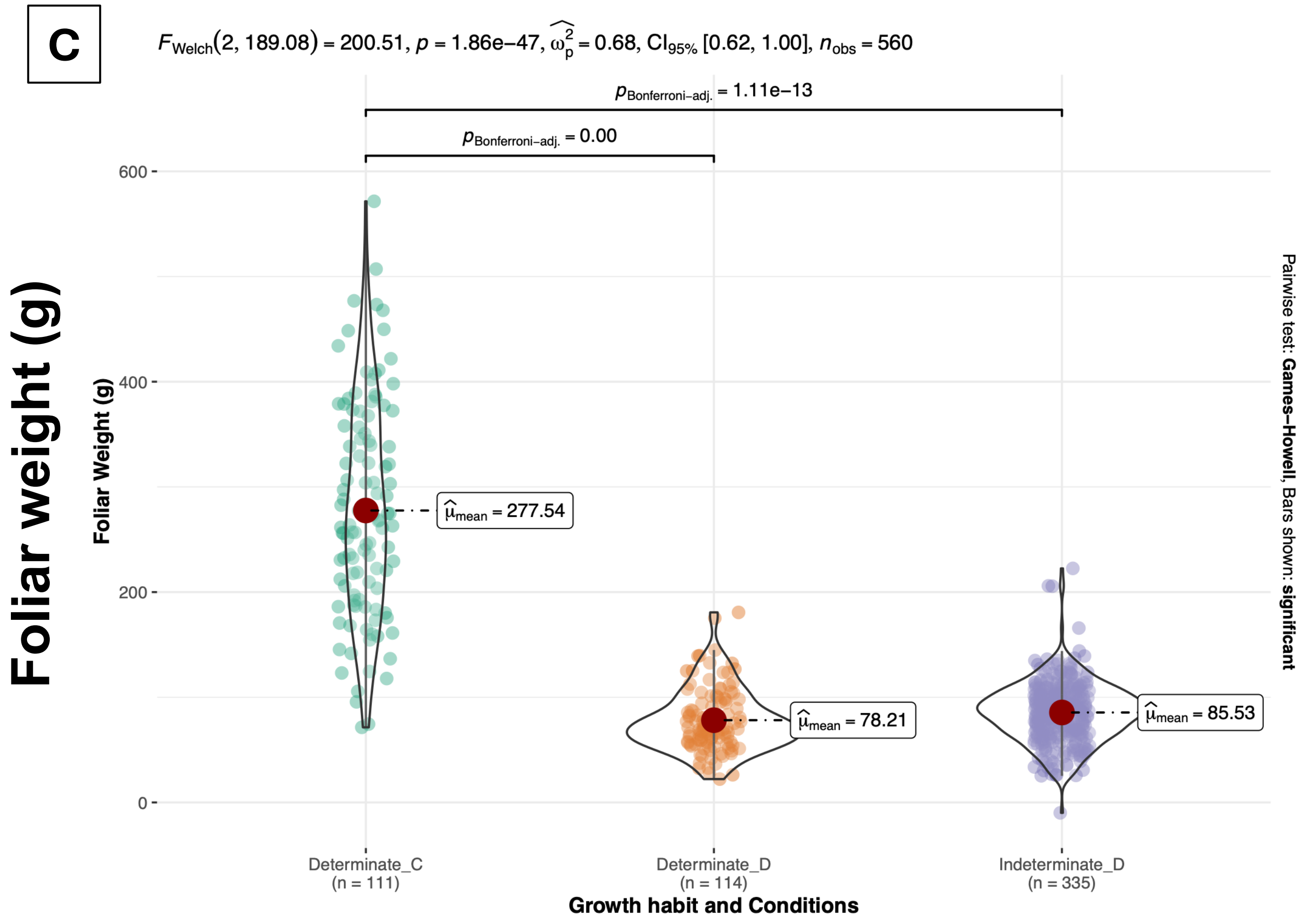

Foliar weight (g)
FWelch(2, 189.08) = 200.51, p = 1.86e−47, ω̂²p = 0.68, CI95% [0.62, 1.00], nobs = 560
pBonferroni−adj. = 1.11e−13
pBonferroni−adj. = 0.00
Foliar Weight (g)
600
400
200
0
μ̂mean = 277.54
μ̂mean = 78.21
μ̂mean = 85.53
Determinate_C
(n = 111)
Determinate_D
(n = 114)
Indeterminate_D
(n = 335)
Growth habit and Conditions
Pairwise test: Games−Howell, Bars shown: significant


D

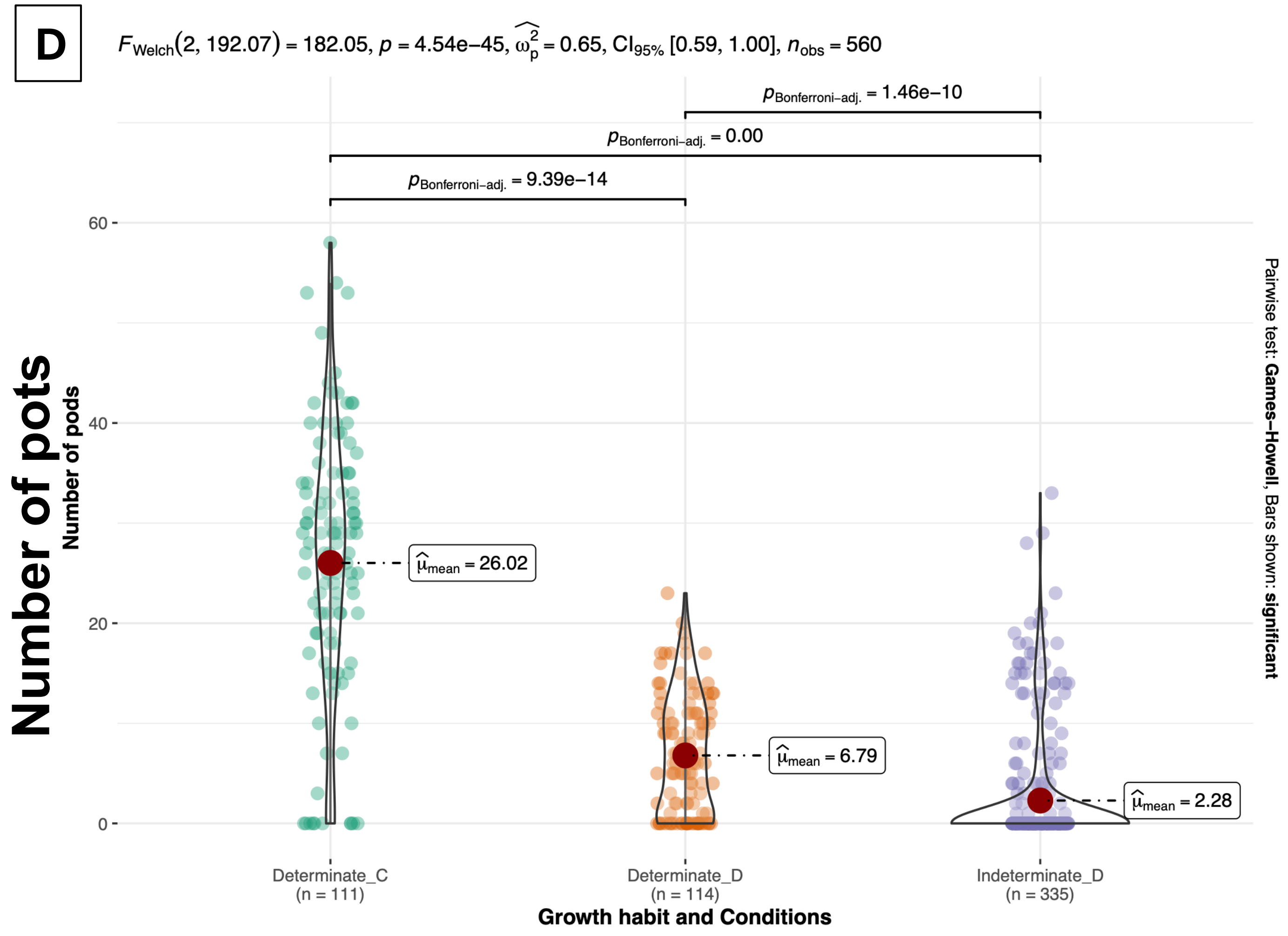

Number of pots
FWelch(2, 192.07) = 182.05, p = 4.54e−45, ω̂²p = 0.65, CI95% [0.59, 1.00], nobs = 560
pBonferroni−adj. = 1.46e−10
pBonferroni−adj. = 0.00
pBonferroni−adj. = 9.39e−14
Number of pods
60
40
20
0
μ̂mean = 26.02
μ̂mean = 6.79
μ̂mean = 2.28
Determinate_C
(n = 111)
Determinate_D
(n = 114)
Indeterminate_D
(n = 335)
Growth habit and Conditions
Pairwise test: Games−Howell, Bars shown: significant
