## supplementary figure s8 for "Developmental and physiological profiles define drought response diversity and genomic associations in common bean"

Suppl fig S8: Comparison of leaf porometer and fluorometer responses per accession (n=142) between water deficit (Week 4) and recovery (Week 5). Significance calculated with Wilcoxen signed-rank test for non-parametric paired data. (NS; not significant, * p <0.05, ** p<0.01, *** p<0.001).

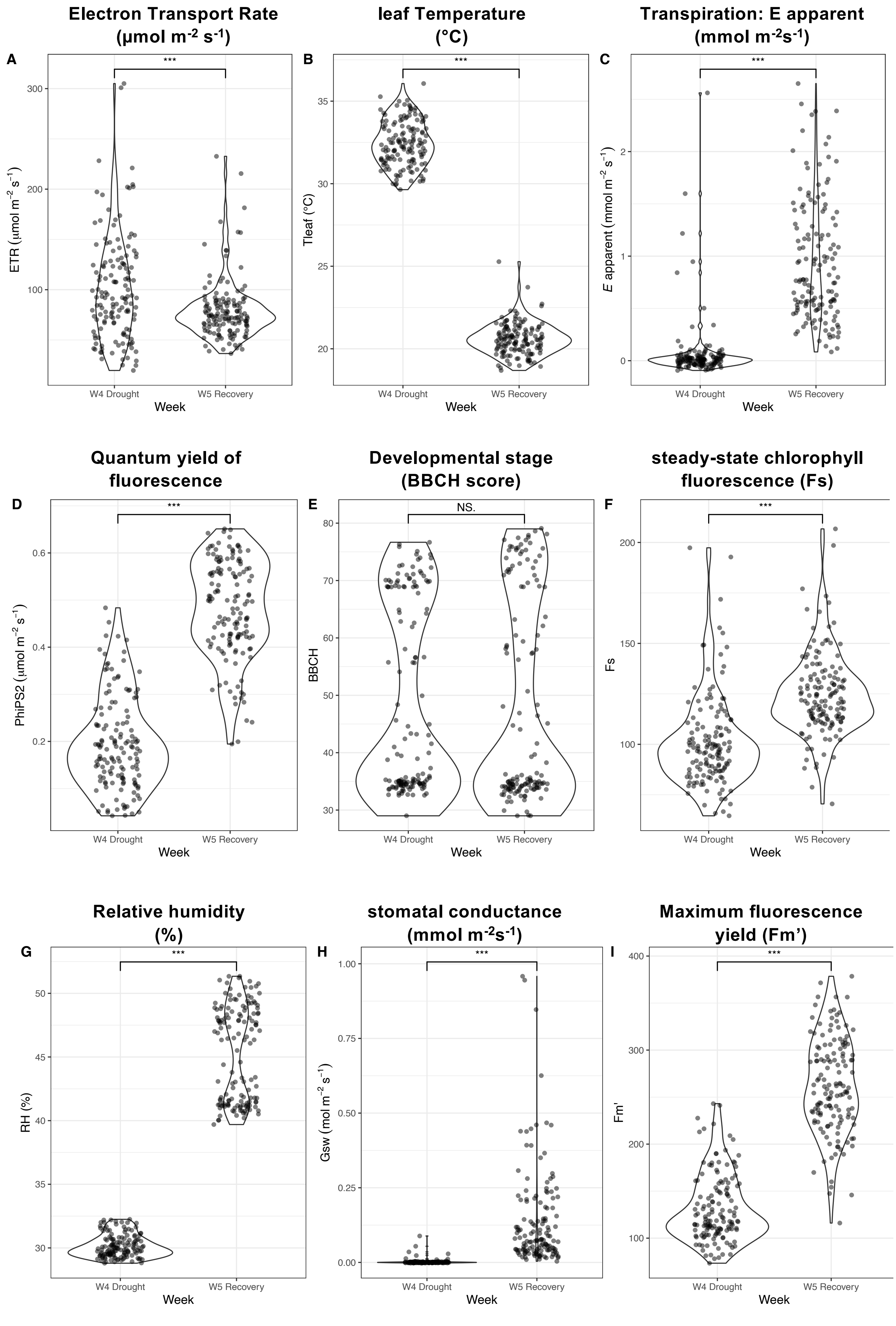
