## supplementary figure s9 for "Developmental and physiological profiles define drought response diversity and genomic associations in common bean"

Supplementary figure S9: Quantile-quantile (QQ) plots per model and grouped by related traits, observed vs expected –log10(p-value) for the association with each phenotype/trait.

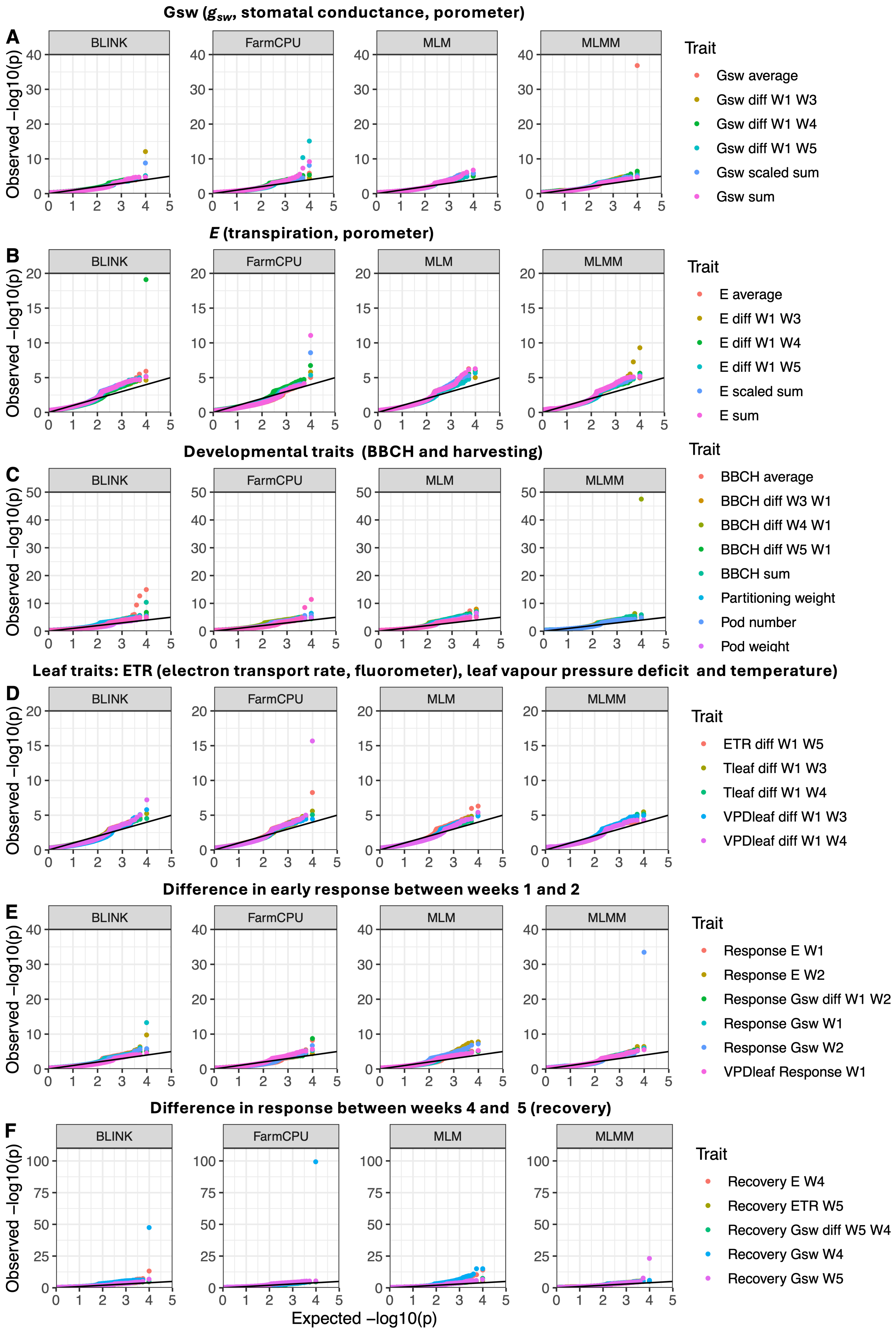
